# Recruiting RyRs to open in a Ca^2+^ release unit: Single-RyR gating properties make RyR group dynamics

**DOI:** 10.1101/821900

**Authors:** D. Gillespie

## Abstract

In cardiac myocytes, clusters of type-2 ryanodine receptors (RyR2s) release Ca^2+^ from the sarcoplasmic reticulum (SR) via a positive feedback mechanism where fluxed Ca^2+^ activates nearby RyRs. While the general principles of this are understood, less is known about how single-RyR gating properties define the RyR group dynamics in an array of many channels. Here, we examine this using simulations with three models of RyR gating that have identical open probabilities. The commonly-used two-state Markov gating model produces frequent, large, and long Ca^2+^ release events because the single exponential that defines its open time (OT) and closed time (CT) distributions reproduces the experimental data poorly. In contrast, simulations that utilize complete single-channel OT and CT distributions fit with multiple exponentials produce infrequent Ca^2+^ release events with far fewer open RyRs. Moreover, when experimentally-measured correlations between single-channel OTs and CTs are included, Ca^2+^ release events become even smaller. This occurs because the correlations produce a small but consistent bias against recruiting more RyRs to open during the middle of a Ca^2+^ release event, between the initiation and termination phases (which are unaltered compared to the uncorrelated simulations). Beyond the effects of full OT and CT distributions and OT/CT correlations on SR Ca^2+^ release, we also show that Ca^2+^ release events can terminate spontaneously without any reduction in SR [Ca^2+^] or physical coupling between RyRs when Ca^2+^ flux is below a threshold value. This both supports and extends the pernicious attrition/induction decay hypothesis that SR Ca^2+^ release events terminate below a threshold Ca^2+^ flux.

**STATEMENT OF SIGNIFICANCE:** This work provides insights into RyR2-mediated Ca^2+^ release by a cluster of RyRs interacting only via their fluxed Ca^2+^. It is shown that: 1) common proxies like the single-RyR open probability versus cytosolic [Ca^2+^] curve and mean open or closed times are poor predictors of SR Ca^2+^ release dynamics; 2) Ca^2+^ release events can self-terminate below a flux threshold without any physical coupling between channels; 3) commonly-used two-state Markov gating models can produce qualitatively different Ca^2+^ release events (larger and longer) compared to simulations where complete single-channel open and closed time distributions are used; 4) correlations between a RyR’s open times and previous closed duration (and vice versa) significantly limit Ca^2+^ release by tamping down the number of open RyRs.

## INTRODUCTION

Over the last two decades, modeling of Ca^2+^ release from the cardiac sarcoplasmic reticulum (SR) through Ca^2+^ release units (CRUs) has revealed the underlying nature of Ca^2+^ sparks, calcium-induced calcium release (CICR), and intracellular Ca^2+^ movement and cycling during a heart muscle contraction (1-14). At the core of these macroscopic processes is a fundamental nanoscale process, the release of Ca^2+^ through ryanodine receptors (RyRs) in the membrane of the SR. RyRs are both Ca^2+^-conducting and Ca^2+^-activated channels that release Ca^2+^ from the SR via a positive-feedback system: an initial Ca^2+^ flux (either through dihydropyridine receptors or a random RyR opening) activates one or more RyRs (CICR) whose fluxed Ca^2+^ activates more and more RyRs (inter-RyR CICR) within the cluster of RyRs that defines a CRU. While much is known about the downstream effects of this Ca^2+^ release (e.g., the making of a spark), relatively few details are known about the mechanism of inter-RyR CICR.

Because there are no experiments that can directly measure what occurs in a CRU, simulations are the only avenue to explore the nanoscale dynamics of RyRs opening and closing due to fluxed Ca^2+^. There are two main computational difficulties. First, it is difficult to make Markov gating models from single-RyR data that elucidate all the open and closed states (15,16), especially under physiological ionic conditions where open probability (*P*o) is low and currents small. Second, modeling Ca^2+^ flux and its buffering by various chelators in the very narrow subsarcolemmal space (∼10–15 nm tall and ∼400 nm wide) is computationally extremely challenging. To sidestep these challenges, it is common to use simplified gating models like two-state gating models with one open (conducting) and one closed (nonconducting) state (C↔O) (2-9,17) defined by single-RyR mean open times (MOTs) and mean closed times (MCTs) (18). Also, the Ca^2+^ movement in the subsarcolemmal space is often simplified, for example by assuming the [Ca^2+^] to be homogeneous throughout the subspace and other compartments (6-11).

These approximations have been vital because they make calculations of Ca^2+^ sparks and intracellular Ca^2+^ movement possible, and therefore are directly responsible for our basic understanding of these processes. With this foundational knowledge in hand, we must now develop a more nuanced understanding of RyR group dynamics within a CRU RyR cluster. Only then will we be able to define the pathological effects on RyR-mediated Ca^2+^ release of proarrhythmic mutations (19-22) and diseases like heart failure and atrial fibrillation (23,24), as well as the therapeutic effects of RyR-targeted drugs and drug candidates (25-27). These mutations, diseases, and drugs, as well as regulatory proteins, alter RyR gating and Ca^2+^ sensitivity. Therefore, we must first understand how single-RyR gating properties affect the group dynamics of a multi-RyR cluster. This is our goal here.

There are several open questions in particular that we want to probe in this study:

1. Are composite quantities that describe RyR gating like *P*o sufficient to give at least a qualitative prediction of SR Ca^2+^ release? If not, are MOT and MCT, the constitutive parts of *P*o, sufficient? These quantities are commonly used to assess the “Ca^2+^ sensitivity” of the RyR, but here we find that none of these are enough to give even a qualitative assessment of Ca^2+^ release; the entire distributions of open times (OTs) and closed times (CTs) is required, not just their mean values. This has ramifications for modeling of SR Ca^2+^ release, as most simulations currently rely on gating models derived solely from MOTs and MCTs.
2. What is the impact of correlations between RyR OTs and CTs and what might their physiological role be, if any? We recently showed that RyR OTs are highly correlated to the length of previous closure, and vice versa (28). These can be seen in Fig. S3. Moreover, we showed that RyR only responds to cytosolic Ca^2+^ when it is closed. These observations have not been included in previous simulations, and here we probe the importance of the OT/CT correlations. We find that the correlations have a moderating effect on SR Ca^2+^ release, making release events with fewer open RyRs. (The lack of Ca^2+^ response in the open state prevents RyR from activating itself with its own fluxed Ca^2+^. Since this is not seen in experiments, we do not explore it further.)
3. Can RyRs in a cluster that interact only via their fluxed Ca^2+^ initiate and terminate release? We examine what such clusters can and cannot do in the absence of things like physical coupling between RyRs, calmodulin, calsequestrin, and low luminal [Ca^2+^]. These are gating modulators or have been proposed as Ca^2+^ release termination mechanisms. By stripping away these factors, one can start to define their roles and assess whether they are necessary for termination or act as modulators. Previous work by us and others has given rise to the idea that termination occurs automatically and inevitably in clusters of RyRs that interact only via fluxed Ca^2+^. Specifically, there is a threshold Ca^2+^ flux below which RyR interactions are not strong enough to sustain release (a mechanism named pernicious attrition by us (13) and induction decay by others (4,5)). Since this mechanism is similar to a phase change seen in statistical mechanics (29,30), it should be unavoidable. Here, we find further support for this idea, while future work will focus on the specific effects of calsequestrin and luminal [Ca^2+^] on regulating this termination mechanism.

We probe these questions by using a very simple model of inter-RyR CICR. The idea is that, with a stripped-down nonphysiological system, we can elucidate physiologically-relevant information about RyR clusters in general. This approach is akin to single-RyR recordings in a bilayer; the system is far from that in vivo, but historically they have revealed physiologically relevant properties like RyR’s Ca^2+^ responsiveness and what factors modulate gating, selectivity, and conduction.

In our reduced model we do not model the geometry of the subsarcolemmal space, cytosolic Ca^2+^ buffers, or depletion of Ca^2+^ from the SR. Rather, each RyR is a point source of Ca^2+^ whose flux is constant when it is open (but the channel may open and close repeatedly) with Ca^2+^ diffusing radially outward into an infinitely large reservoir. It is as if we placed an array of RyRs into the bilayer of a single-channel experiment and recorded the resulting openings and closings. Since the RyRs still interact via Ca^2+^, we will be able to see what factors significantly alter the RyRs’ response and group dynamics. In this way, our results will provide physiological insights because our underlying single-RyR data was taken under physiologically relevant conditions and because the RyR group dynamics seen in this reduced system are likely present in more complex systems.

To address the questions listed above with this system, we consider three subtly different models of RyR gating and compare the results. The three models of single-RyR gating are all derived from the same experimental data, and at every measured cytosolic [Ca^2+^] the three models have identical MOT, MCT, and *P*o. They differ in how their OT and CT distributions are summarized for each cytosolic [Ca^2+^]: one model is the equivalent of a two-state Markov gating model and fits a single exponential distribution to the OTs and CTs; another model fits them with up to 5 exponentials; the third model also uses multiple exponentials, but as described later, takes into account the measured correlations between OTs and CTs. As a shorthand, we refer to these as the two-state, uncorrelated multi-*τ*, and correlated multi-*τ* gating schemes, respectively. (Multi-*τ* refers to the multiple time constants *τ*, as opposed to the single time constant for the two-state model.)

Collectively, our simple model reveals several new insights into RyR2 group dynamics in a CRU and SR Ca^2+^ release in general:

1. The single-RyR *P*_o_ versus cytosolic [Ca^2+^] curve is a poor predictor of SR Ca^2+^ release properties. By design, all three of our gating models have identical *P*_o_ curves, but all three have very different release properties, especially for the largest release events. Moreover, SR Ca^2+^ release was qualitatively different just by switching from the two-state gating scheme to the uncorrelated multi-*τ* scheme. The only difference between the two is how many exponentials are used to describe the OT and CT distributions. Thus, we conclude that RyR group dynamics depend strongly on the details of the single-RyRs gating properties and how well the gating model recapitulates the single-RyR OT and CT distributions.
2. The correlations between RyR OTs and CTs significantly decrease the number of RyRs open during a Ca^2+^ release event (which we define throughout as first channel open to last channel closed). Interestingly, they do not alter release event initiation or termination, but instead only limit the recruitment of more channels to open. This core phase of Ca^2+^ release occurs between initiation (the initial recruitment of a few channels) and termination (the closing of the last few channels) and determines the number of RyRs open during the main part of the release event. To our knowledge, this phase of Ca^2+^ release has not been considered in detail before.
3. In the model, the RyRs interact only via their fluxed Ca^2+^, and if the flux is sufficiently small, the Ca^2+^ release events terminate spontaneously even though there is no SR Ca^2+^ depletion; a decrease in SR [Ca^2+^] is not necessary to stop release. This lends support to the pernicious attrition idea that there is a threshold Ca^2+^ flux below which release events cease (4,5,13). Those works assume that a decreasing SR [Ca^2+^] is vital for termination. Our results, with a constant SR [Ca^2+^], extend the pernicious attrition hypothesis by showing SR depletion is not strictly necessary for Ca^2+^ release self-termination; there is a threshold of constant Ca^2+^ flux below which Ca^2+^ release events always terminate. (While Stern’s stochastic attrition (31) describes termination with constant Ca^2+^ flux, the idea of a threshold Ca^2+^ flux was, to our knowledge, not described by Stern.) Overall, this work supports the idea that termination is a built-in property of RyRs in clusters and that molecular modulators of gating potentially regulate that process but may not drive it (but more on this needs to be done).
4. The two-state gating scheme, which has been used in many earlier studies (2-9,17), behaves qualitative differently during all three phases of Ca^2+^ release from our other gating schemes (the two multi-*τ* schemes). If initiation of a release event is defined as the opening the first few channels (e.g., 3), then initiation is the rate at which 1- and 2-channel events do not end before they become 3-channel events. The two-state scheme is significantly more successful at converting 1- and 2-channel events into release events with 2 and 3 channels opens, respectively; it is far more efficient at initiating a Ca^2+^ release event than the multi-*τ* schemes. During the core phase of the Ca^2+^ release, the two-state scheme is similarly more successful at opening other RyRs than either multi-*τ* scheme, leading to more open RyRs per event. Lastly, the time required to close the last few RyRs is much shorter for the two-state scheme. This suggests that two-state models are likely not the best choice to replicate how RyRs work in clusters. We propose that this is because the two-state model is a poor representation of the single-RyR OT and CT distributions. With its single-exponential for these distributions, it severely underrepresents both short and long events in the single-channel distributions and therefore in the simulations.

Lastly we note one aspect of our simulation method that may be of wider interest to the modeling community. Specifically, we use an algorithm to stochastically flip channel states that may be an effective alternative to traditional Markov gating models. It is fast and easy to implement because it directly translates the OT and CT distributions into closing and opening probabilities after fitting the data to a sum of exponentials (see Supporting Material). As such, it does not provide physical insights into the gating process like a Markov model might; increasing the number of exponentials has no physical interpretation in our algorithm. However, our algorithm is much easier to set up because the sole purpose of our fitting is to accurately reproduce the data, where increasing the number of exponentials can be helpful.

## THEORY AND METHODS

We briefly describe the experiments, the simulations, and the three gating schemes, with complete details given in the Supporting Material.

### Experiments

The single-RyR data we use has been previously published (32) and analyzed (28). Specifically, rat RyR2s were recorded in artificial bilayers with 1000 µM luminal (intra-SR) Ca^2+^ and with 0.1, 1, 10, 50, 200, and 1000 µM cytosolic Ca^2+^ with cell-like cytosolic Mg^2+^ and ATP levels, which are potent modulators of RyR2 gating (33,34). This ensures gating and divalent ion concentrations are as close to physiological as possible, while a large Cs^+^ gradient was used to make large, easily measured currents. Further details are in the Supporting Material and Tables S1 and S2.

As described in Ref. (28), our native (non-purified) RyRs exhibit different modes of gating. RyRs in cells are subject to post-translational modifications like phosphorylation and oxidation and are also known to be associated with various protein partners like calmodulin, calsequestrin, FKBP, kinases, junctin, and triadin. The single-RyR recordings used here were made by fusing native SR microsomes into planar lipid bilayers. Consequently, the post-translational modification status and/or protein compliment of the RyRs fused into our planar bilayers is likely varies channel to channel, not unlike the situation within living cells.

In our simulations here, we use an averaged version of the different modes, lumping all the data together so produce OT and CT distributions. This does not affect our results, as our aim is not complete physiological accuracy, but rather to understand how changes in the descriptions of single-RyR gating (e.g., using one exponential or multiple exponentials for the OT and CT distributions) affect Ca^2+^ release through multiple RyRs in an array.

### Simulations

#### A simple geometry

We do not model the geometry of the subsarcolemmal space, cytosolic Ca^2+^ buffers, or depletion of Ca^2+^ from the SR. Rather, each RyR is a point source of Ca^2+^ whose flux is constant when it is open (the channel may open and close repeatedly) with Ca^2+^ diffusing radially outward into an infinitely large reservoir. It is as if we placed an array of RyRs into the bilayer of a single-channel experiment.

This approach has pros and cons. The biggest drawback is lack of in situ realism (e.g., no confining geometry, no Ca^2+^ buffers). On the other hand, working in a simpler geometry facilitates studying RyR group dynamics (the goal of this work) by removing complicating factors to focus purely on the factors affecting Ca^2+^-activated release. The idea is that our results will provide physiological insights because our underlying single-RyR data was taken under physiologically relevant conditions and because RyR group dynamics seen in a simple system are likely present in more complex systems. Overall, RyRs react to Ca^2+^, whether it was buffered first or not, and our approach captures this; buffers modulate [Ca^2+^] but do not change the underlying actions of Ca^2+^ on RyR. The buffering, especially the fast buffering of the sarcolemmal membranes (35,36), will affect the details of RyR group dynamics, but not our conclusions about single-RyR gating properties defining Ca^2+^ release events.

From a numerical point of view, our simplified geometry and lack of Ca^2+^ buffers is a computationally-tractable way to compute sufficient numbers of rare Ca^2+^ release events to be statistically relevant. Currently, a full 3D reaction-diffusion system with subnanometer and submicrosecond spatial and temporal resolution that can run minutes of simulated time is not practical. However, a recently derived analytic solution to the spherically-symmetric diffusion equation from a point source with variable flux (37) (see Supporting Materials) makes our simulations fast enough to compute hours of simulated time and millions of Ca^2+^ release events.

Moreover, the use of a constant Ca^2+^ flux is useful for our analysis. By maintaining the same current all the time when channels are open, during a Ca^2+^ release event the cytosolic [Ca^2+^] seen by closed RyRs remains consistent. Specifically, if, for example, 4 RyRs are open, then the other closed RyRs are exposed to a cytosolic [Ca^2+^] distribution (Fig. S2). This is same whenever 4 RyRs are open, regardless of the gating scheme used or RyR array size (data not shown). Therefore, any differences in RyR group dynamics are not due to differences in cytosolic [Ca^2+^] felt by other channels in the array, as might happen with a variable SR [Ca^2+^].

Lastly, by stochastically gating our channels by sampling the OT and CT distributions (see below), rather than using a Markov gating model, we can directly test the importance of the RyR OT/CT correlations on RyR group dynamics by simply turning off any memory of the previous state.

#### Stochastic gating

All RyRs are closed at the beginning of the simulation. When one RyR opens randomly, Ca^2+^ diffuses to nearby RyRs that may react by opening, and flowing more Ca^2+^ into the system. One crucial aspect then is how RyRs open and close, both randomly and in response to Ca^2+^. Here, we do not construct traditional Markov gating models from the experimental single-channel data. Instead, our simulations stochastically open and close channels with a probability derived from the distribution of open times (OTs) and closed times (CTs), as described by Colquhoun and Hawkes (18).

As an example, consider an RyR that has been open for some amount of time *T*. The probability that it will close during the next timestep Δ*t* is the conditional probability that it will close between *T* and *T* + Δ*t* given that it has already been open for time *T*. Using the shorthand notation of *o* @ *t* and *c* @ *t* to denote open at time *t* and closed at time *t*, respectively, this conditional probability is (18)

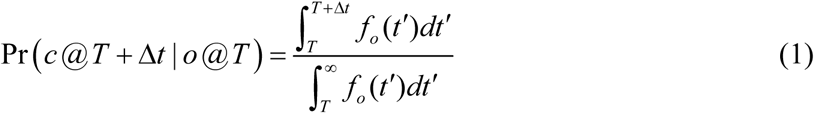

Where *f*_*o*_ (*t*’) is the probability distribution function of the OTs. This function is derived from the experimentally measured distributions of OTs by fitting it with a sum of exponentials (see below). At every timestep in the simulation, Eq. (1) is used to determine whether an open channel closes by drawing random numbers from the probability distribution *f*_*o*_ (*t*’). (A similar probability is used for opening a closed channel, but based on *f*_*c*_ (*t*’), the probability distribution function of CTs.) The *f*_*o*_ (*t*’) and *f*_*c*_ (*t*’) are derived from the data at every experimental cytosolic [Ca^2+^] (as described below for the three gating schemes) and then interpolated to other [Ca^2+^], which is necessary since different RyRs in the array see different cytosolic [Ca^2+^] at any one time.

The open/closed state of each channel in the array is updated at each timestep depending on the [Ca^2+^] each sees, with channels changing state based on randomly-generated numbers and the probability of flipping states defined by Eq. (1).

Detailed descriptions of how the gating was implemented, how the simulations were performed, and how the *f*_*o*_ (*t*’) and *f*_*c*_ (*t*’) were interpolated to non-experimental cytosolic [Ca^2+^] are in the Supporting Materials.

#### Incorporating recent findings

In Ref. (28) it was shown that RyR does not respond to its own fluxed Ca^2+^ because it responds to cytosolic [Ca^2+^] only when it is closed. Moreover, it was shown that the duration of an opening is strongly correlated to the duration of the previous closure, and vice versa that the CT is correlated to the OT of the previous opening. Unlike previous studies, we incorporate these two findings in the simulations. (The technical details of how this is done are described in the Supporting Materials.)

The OT/CT correlations are recapitulated in Fig. S3. Here, one focus in particular is comparing gating schemes with and without this memory of the previous state’s duration. In yet unpublished single-RyR recordings in other species and in the presence of calsequestrin, these correlations are consistently found (personal communication, Michael Fill, Rush University, July 2019), and so we believe them to be physiologically relevant.

### Three RyR gating schemes

The most common way to model gating of channels is via Markov models, with a number of so-called open and closed states and one conducting state. While Markov models with multiple states better represent the single-RyR OT and CT data, it is difficult to elucidate all the open and closed states (15,16). Therefore, two-state Markov RyR gating models are very common in simulations of SR Ca^2+^ release (2-9,17).

In a two-state gating scheme, the channel switches between a conducting and nonconducting state, and the distributions of OTs and CTs are a single-exponential fit of the data with the time constant being the measured MOT and MCT (18). This is one of the gating schemes we will use here, where the *f*_*o*_ (*t*’) (and *f*_*c*_ (*t*’)) in Eq. (1) is this single-exponential function. Colquhoun and Hawkes (18) showed that a two-state Markov gating model is equivalent to using Eq. (1) in a simulation.

By definition, this gating scheme recapitulates the single-RyR MOT and MCT for each cytosolic [Ca^2+^] and therefore the experimental *P*_o_ versus cytosolic [Ca^2+^] curve (black symbols in Fig. 1A). However, the single-exponential *f*_*o*_ (*t*’) and *f*_*c*_ (*t*’) OT and CT distribution functions do a poor job of reproducing the single-RyR data (black curves in Figs. 1A and 1B, respectively). In the simulation, it is these probability density functions that are sampled and therefore the two-state model will have few short and long openings and closings since the black curves in Figs. 1A and 1B miss these almost entirely. (Note that adjusting the two-state to include missed recorded events does not change this, as shown in the Supporting Materials.)

**Fig 1.**
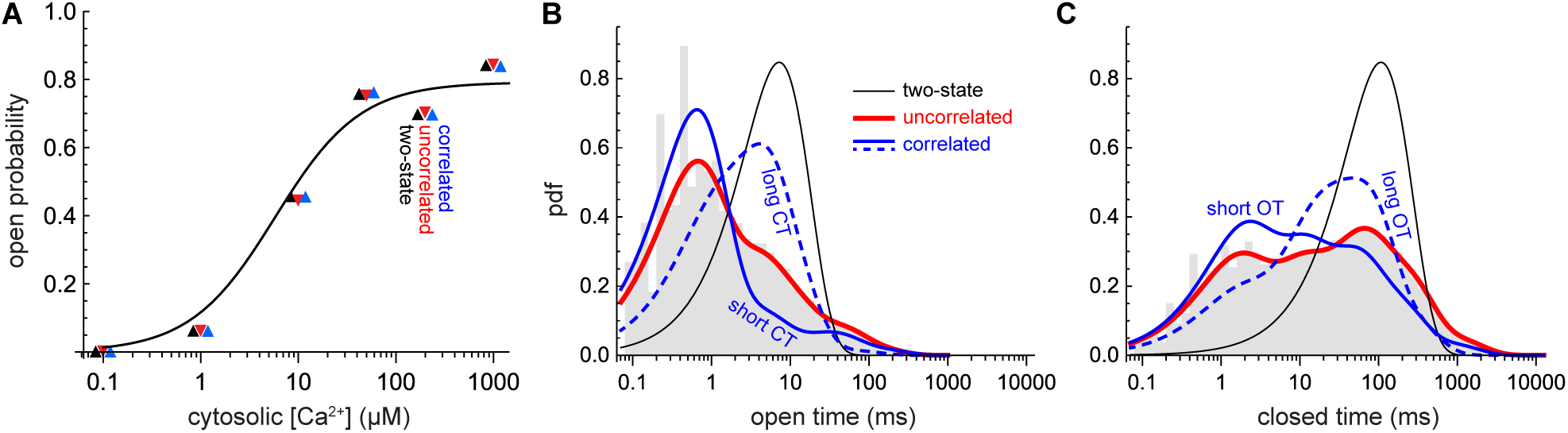
(color online) (A) *P*_o_ versus cytosolic [Ca^2+^]. The symbols are for the three gating schemes with a coloring scheme used throughout (black: two-state; red: uncorrelated multi-*τ* scheme; blue: correlated multi-*τ* scheme). Note the symbols are offset on the *x*-axis to show they have the same *P*_o_. The solid line is the Hill equation fit of the data taken directly from Ref. (28). (B) OT distributions. Gray bars: experimental data. Black: single-exponential fit. Red: multi-exponential fit. Blue: Multi-exponential fit of OTs with short CTs (0–0.531 ms) of the previous closure (solid line) and of OTs with long (≥530.9 ms) previous closures (dashed). (C) Same as in panel B, but for CT distributions. Blue: Short previous OTs (solid) are 0–0.266 ms and long previous OTs (dashed) are ≥53.1 ms. In panels B and C, the cytosolic [Ca^2+^] is 1 μM. The fit of the blue curves to the corresponding data is shown in Fig. S1.

One goal of this study is to understand whether this has a significant impact on simulated Ca^2+^ release or whether merely having the MOT and MCT correct is sufficient. Therefore, to faithfully reproduce the RyR data, we fitted the OT and CT distributions with up to 5 exponentials. The red curves in Figs. 1B and 1C are the resulting *f*_*o*_ (*t*’) and *f*_*c*_ (*t*’). When fitting the data with multiple time constants *τ*, the fit was again constrained to reproduce the experimental MOTs and MCTs so the *P*_o_ was the same as for the two-state scheme (Fig. 1A, black and red symbols). Sampling these multi-*τ* fitted distributions in our simulations evolves the system via what we call the uncorrelated multi-*τ* gating scheme.

It is important to note that neither of the multi-*τ* schemes is a Markov gating model. We have not defined multiple open and closed states and the number of exponentials used to fit the data have no physical interpretation as they do in a Markov model. In fact, we have explicitly avoided multi-state Markov models because they are difficult, cumbersome, and time consuming to create. By directly using the red curves in Figs. 1B and 1C (the *f*_*o*_ (*t*’) and *f*_*c*_ (*t*’)) in Eq. (1) to define whether a channel switches states from open to closed (or vice versa) during the simulation, we can sidestep the Markov model creation process entirely. In fact, using Eq. (1) to propagate the simulation in time allows us move straight from the experimental data to doing simulations. All that is required is to fit OT and CT distribution data to produce *f*_*o*_ (*t*’) and *f*_*c*_ (*t*’) that accurate reproduce the data.

We call it the “uncorrelated” gating scheme because neither it nor the two-state scheme produce correlations between the OTs and CTs. To flip from open to closed, for example, in these schemes, the complete OT distribution is sampled at each timestep, without any knowledge of the previous closure. To include this memory and produce OT/CT correlations, we break the experimental OT and CT data into smaller parts. For OTs, for example, first all OTs are paired with the CT of the preceding closed state. These pairs are then binned into ranges of CTs so that there are at least 1000 pairs in each bin. The corresponding OTs in each CT bin are then fit with exponentials (up to 5) to produce an OT distribution for each CT bin (blue lines in Figs. 1B and 1C; see also Fig. S1). As before, the fits are constrained to reproduce the experimental mean value of the OTs in each data subset. A similar thing is done to link CTs paired with previous OTs. In a simulation with only one channel, this reproduces the experimental MOTs and MCTs of the full data set and therefore the *P*_o_, as shown in Fig. 1A (blue symbols). Moreover, it reproduces the experimental correlations (Fig. S3). Therefore, we call this the correlated multi-*τ* gating scheme.

In this way we have three differently gated kinds of RyRs: two-state, uncorrelated multi-*τ*, and correlated multi-*τ*. For all the experimental cytosolic [Ca^2+^] (0.1, 1, 10, 50, 200, and 1000 µM) they have the same MOT, MCT, and therefore *P*_o_. Therefore, in simulations with only one channel present, each recapitulates the experimental *P*_o_ versus cytosolic [Ca^2+^] curve (symbols in Fig. 1A).

Where they differ is in their single-RyR OT and CT distributions. Specifically, in single-channel simulations the OT and CT distributions each scheme generates is the one used to define the gating scheme (data not shown). For the two-state scheme that is the black line in Fig. 1B. Both multi-*τ* schemes reproduce the red line in Fig. 1B, but the uncorrelated multi-*τ* scheme does not reproduce the experimental OT/CT correlations (not shown) while the correlated multi-*τ* scheme does (Fig. S3). Therefore, the two-state scheme has OT and CT distributions that compare poorly to the experimental ones (while retaining the experimental MOT, MCT, and *P*_o_), while the multi-*τ* schemes compare as best as we can make them. One level deeper, the correlated multi-*τ* scheme also reproduces the experimental OT and CT distributions that take into account the length of the previous event (Fig. S1).

## RESULTS AND DISCUSSION

Here, we examine how these different RyR gating schemes produce different Ca^2+^ release behavior in an array where the RyRs interact only via their fluxed Ca^2+^. By comparing how similar the Ca^2+^ release events are for three gating schemes that retain the same MOT, MCT, and *P*_o_ versus cytosolic [Ca^2+^] curves we assess how well composite quantities like MOT, MCT, and *P*_o_ can predict SR Ca^2+^ release or whether the recapitulation of full OT and CT distributions is necessary. In addition, by comparing the results of the two multi-*τ* schemes, we hope to assess what the functional effect of the OT/CT correlations is and whether they might be physiologically significant. Lastly, by varying the size of the Ca^2+^ flux by which the RyRs interact we hope to get a better understanding of how such a CRU works and under what conditions it can self-terminate Ca^2+^ release events.

At the start of our simulations, all RyRs in the array (ranging from 3×3 to 14×14 in size) are closed and only a random opening perturbs the system, as in diastole, to kick off Ca^2+^ release. Also to mimic diastole, the background cytosolic [Ca^2+^] is 0.1 µM. Representative traces of the number of open channels versus time are shown in Fig. 2A.

**Fig 2.**
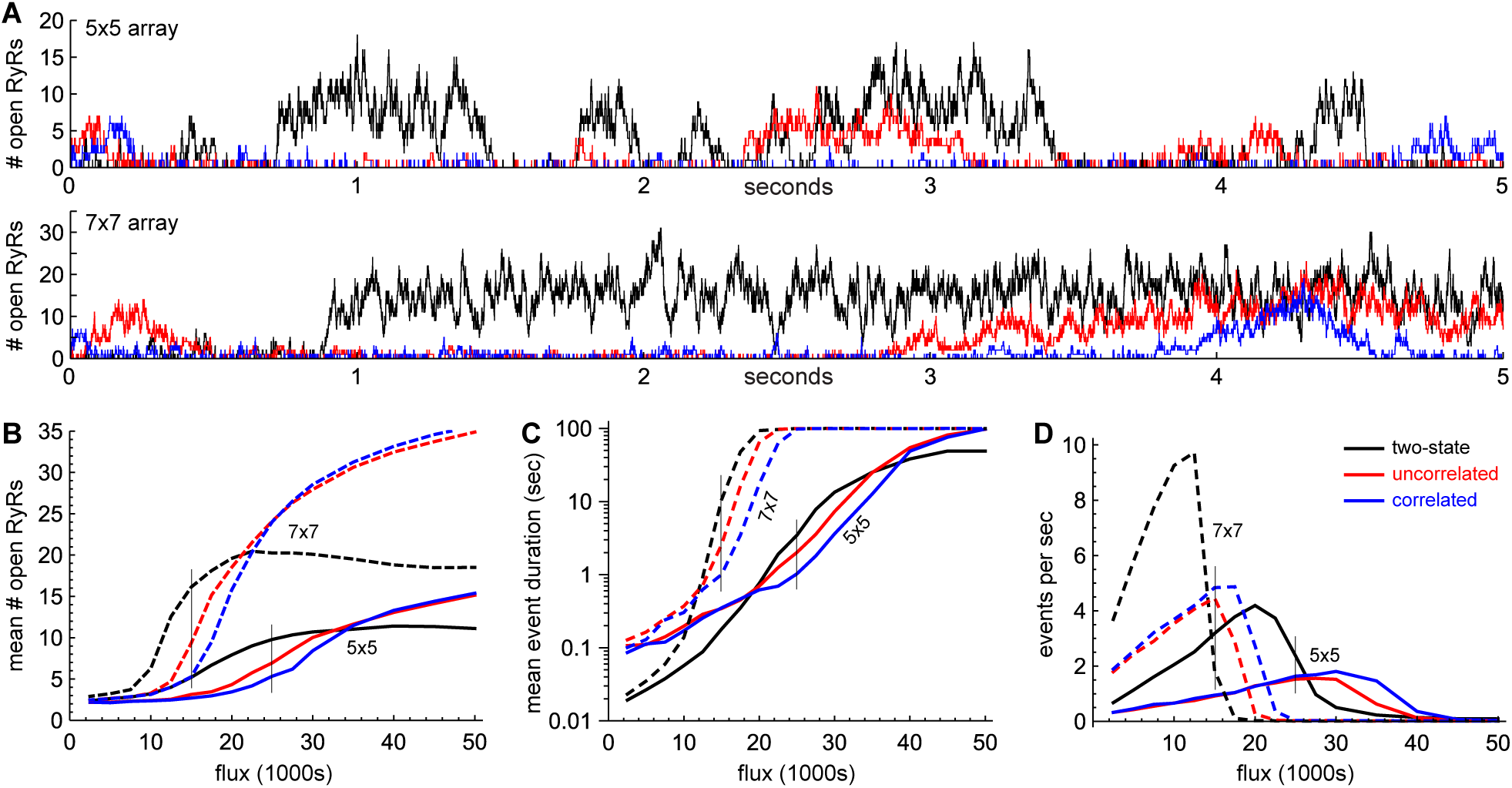
(color online) (A) Timeseries of the number of open RyRs during the first 5 sec of simulations using the two-state scheme (black), uncorrelated multi-*τ* scheme (red), and the correlated multi-*τ* scheme (blue) for a 5×5 (top) and 7×7 (bottom) array for fluxes of 25,000 and 15,000 s^−1^, respectively (vertical lines in panels C–D). (B) Mean number of open RyRs of the largest Ca^2+^ release event in each of 25 100-sec long simulations done for each unitary RyR Ca^2+^ flux for a 5×5 (solid) and 7×7 (dashed) array. Color scheme as in panel A. (C) Mean duration of open RyRs of the longest Ca^2+^ release event in each of the simulations. Color scheme and dashing as in panel B. (D) Frequency of Ca^2+^ release events with at least 3 open RyRs. Color scheme and dashing as in panel B. More detailed frequency data and the equivalent of panels B and C for a 10×10 array are shown in Fig. S4. The vertical lines in panels B–D are the fluxes used in panel A and in the analysis shown in Figs. 3 and S5.

### Termination

We start by noticing that Ca^2+^ release events terminate spontaneously, even though the unitary RyR Ca^2+^ flux is constant in time. This has been reported before (31), but here we show this is true when the flux is small enough, while above a threshold flux (which varies by array size) Ca^2+^ release can continue indefinitely. The threshold fluxes are seen in Figs. 2D and S4 where the event frequencies decrease rapidly with increasing Ca^2+^ flux. Above the threshold, Ca^2+^ release events do not terminate and the frequency of events drops to 1 event for the entire 100-sec long simulation. As array size increases, the interval over which the frequency declines narrows and the transition from many events to one event sharpens; it becomes more of true threshold phenomenon as the array size increase. (As a technical aside, this behavior is consistent with a phase transition, where sharpening of transitions is expected for increasing array size (38).)

Below the threshold Ca^2+^ flux, *all* of the *millions* of Ca^2+^ release events in our simulations terminated spontaneously for all three gating schemes. Even when so many channels are open so that the neighboring closed channels see >10 µM cytosolic [Ca^2+^] (Fig. S2) where the *P*_o_ is >50% (Fig. 1A), every release event eventually stopped for the conditions shown in Fig. 2A. This is consistent with recent work showing there is a flux threshold below which continuous Ca^2+^ release is not possible (4,5,13,29,30). In fact, it strengthens this pernicious attrition/induction decay hypotheses because here we do not rely on a decreasing Ca^2+^ flux to produce or facilitate termination. For large fluxes, some SR Ca^2+^ depletion (which the current model does not include) is likely necessary to decrease the flux sufficiently to end Ca^2+^ release, but this remains to be verified.

Overall, this shows that, under the right conditions, SR Ca^2+^ release termination in a CRU where RyRs interact only via Ca^2+^ is not only possible, but inevitable. However, it still remains to be shown in future work how different RyR array spacings REF and modulators of RyR gating that we have not considered here (e.g., calmodulin, calsequestrin, low SR [Ca^2+^]) will alter termination.

### Different gating schemes, different Ca^2+^ release

Next, we turn to the main analysis involving the three gating model.

#### Two-state gating is qualitatively different

It is evident from the traces in Figs. 2A that the three gating schemes have very different group dynamics. As discussed in detail below, the two-state scheme (black lines) produces Ca^2+^ release events that are longer and have more open channels than the two multi-*τ* schemes. In turn, the uncorrelated multi-*τ* scheme (red) has more open channels per release event than the correlated multi-*τ* scheme (blue).

As shown in Fig. 1A, all three gating schemes produce identical *P*_o_ versus cytosolic [Ca^2+^] curves because, by design, each fitting with exponentials of OT and C distributions was constrained to reproduce the experimental MOTs and MCTs. Therefore, any differences in how the channels behave in a cluster of RyRs are not due to different channel Ca^2+^ affinities (i.e., shifts of the *P*_o_ curve to the left or right) or different MOTs and MCTs that retained the same *P*_o_ (e.g., the MOTs and MCTs were not both 50% smaller in one gating scheme than in another).

A notable difference between the three gating schemes is in the large release events (i.e., the Ca^2+^ release events with a large average number of channels open). To illustrate this, we performed 25 simulations lasting 100 sec for different unitary RyR fluxes. The means of the largest and longest-lasting events from each simulation are shown in Fig. 2B and 2C for 5×5 and 7×7 arrays, respectively. (10×10 arrays are shown in Fig. S4.) The two-state scheme (black lines) behaves significantly differently than either of the multi-*τ* schemes. At low unitary RyR flux, the large two-state events have far more channels open than the multi-*τ* schemes’ release events, and the longest events tend to be much shorter. At high flux, however, the exact opposite is true. Moreover, the frequency of large Ca^2+^ release events is qualitatively different; the two-state scheme produces a higher rate of events with multiple channels open (Fig. 2D) and a lower rate single-channel events compared to either multi-*τ* scheme (Fig. S4).

One important result seen in Figs. 2 and S4 is the substantial differences between the two-state and uncorrelated multi-*τ* schemes (black and red curves, respectively). In the simulations, the only difference between them is how many exponentials are used to fit the OT and CT distributions (Figs. 1B and C). Just by moving to a multi-exponential fit (red lines) of the same data but otherwise changing nothing, the group dynamics have changed qualitatively. Since the multi-*τ* fit reproduces the experimental data more faithfully, this indicates that a two-state gating scheme does not seem to capture how RyRs function collectively in a cluster. Such a fundamental difference will also affect Ca^2+^ release in more realistic models of a CRU.

In some sense this is not surprising. From a Markov gating scheme point of view, having only one conducting state and one nonconducting state is convenient and reasonable; the average open and closed times are used to approximate the behavior of the channel. However, looking at this from the point of view of the OT and CT data, the reason for the differences between the two-state and multi-*τ* gating schemes is clear: a single exponential is an extremely poor representation of the data (Figs. 1B and 1C). Even though the black curves and red curves have the same MOT and MCT (Figs. 1B and 1C, respectively), both short and long events are missing in the two-state scheme.

#### Effect of OT/CT correlations

Next, we compare the uncorrelated and correlated multi-*τ* schemes (red and blue curves, respectively, in Figs. 2 and S4). These also exhibit important differences, but to a lesser extent. At low and high unitary RyR Ca^2+^ fluxes, the two behave very similarly, but at intermediate fluxes (shown by thin vertical lines in Figs. 2B–D) the correlated multi-*τ* scheme has fewer number of RyRs open during the largest events (Fig. 2B) and the longest events are shorter (Fig. 2C). Interestingly, both multi-*τ* schemes can have the same frequency of release events with 3 or more RyRs open (Fig. 2D), as well as the same frequency of events with 1 or 2 RyRs open (Fig. S4).

To explore these differences, we grouped Ca^2+^ release events according to the maximum number of open RyRs (*N*_max_). We then computed the fraction of release events for each possible *N*_max_ (i.e., what percentage of release events had a maximum of 1 RyR open, a maximum of 2 open, etc.). This is shown in Fig. 3A for all three gating schemes for the 5×5 array of RyRs. The same is shown in Fig. S5 for the 7×7 and 10×10 arrays. Since events with a large *N*_max_ are rare, we performed 500 100-sec-long simulations at one RyR unitary Ca^2+^ flux (thin vertical lines in Figs. 2B–D) to ensure statistical confidence. In total, the red and blue curves in Fig. 3 each summarize 2.8 million release events, 4.7 million in Fig. S5A, and 6 million in Fig. S5B.

**Fig 3.**
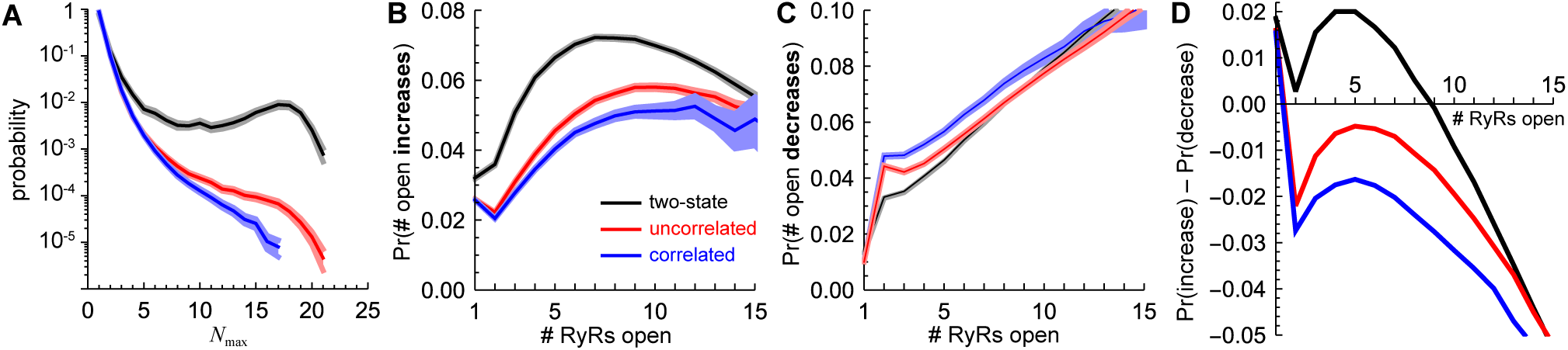
(color online) Release event statistics for a 5×5 array with a unitary RyR Ca^2+^ flux of 25,000 s^− 1^. Black: two-state scheme. Red: uncorrelated multi-*τ* scheme. Blue: correlated multi-*τ* scheme. (A) Probability of having a Ca^2+^ release event for a given *N*_max_, the maximum number of open RyRs during a release event, with a minimum of 10 Ca^2+^ release events required at each *N*_max_. (B) Probability of increasing the number of open RyRs by 1 at the next timestep during a release event when *n* RyRs are currently open (*x*-axis). A minimum of 100 events were required for each *n*. (C) Same as panel B, except is the probability of decreasing the number of open RyRs by 1. (D) Difference of the corresponding solid lines in panels B and C. In panels A–C, the shaded areas are 95% confidence intervals obtained by bootstrap resampling. Specifically, 500 100-sec long simulations were performed and 1000 random samples with replacement of whole simulations were re-analyzed and the 2.5% and 97.5% quantiles are the limits of the confidence intervals (28). The solid line is the quantity obtained from the original (not resampled) data.

The figures show that both the uncorrelated and correlated multi-*τ* schemes have a similar probability of small events (i.e., with *N*_max_ small), but they diverge as *N*_max_ grows. The correlated multi-*τ* scheme produces far fewer large release events than the uncorrelated multi-*τ* scheme. This is significant because large Ca^2+^ release events can be physiologically detrimental. And while ours is not the physiological case, there is no a priori reason to believe that such a fundamental difference would disappear in vivo.

This could reflect a difference in event initiation (i.e., converting a small event with few open RyRs into a big event with many RyRs open). This, however, does not seem to be the case. First, at the flux used in Fig. 3, both multi-*τ* schemes have identical event frequencies for *N* _max_ = 1, *N* _max_ = 2, and for *N*_max_ ≥ 3 (Fig. S4). Specifically, if the uncorrelated multi-*τ* scheme was more successful at recruiting RyRs to open during the initial stages of a release event when the number of open RyRs was small (e.g., 1 or 2), then one might expect lower event frequencies for *N* _max_ = 1 and for *N* _max_ = 2 and higher frequencies for *N*_max_ ≥ 3. But this is not the case (Fig. S4). Second, we directly computed the rate at which 1- and 2-channel events were converted to 3+-events and the rate at which the small events were snuffed out (i.e., self-terminated). These were identical for both multi-*τ* gating schemes: in the 5×5 array for a flux of 25,000 s^−1^, 97.2% of 1- and 2-channel events were snuffed out and only 2.8% grew to 3+-channel events; in the 7×7 array for a flux of 15,000 s^−1^, 95% small events terminated and 5% grew larger. (The two-state gating scheme is qualitatively different in another way. For the 5×5 array, it had a conversion rate of ∼14%, 3 to 5 times that of the multi-*τ* schemes; only ∼86% of 1- and 2-channel events were snuffed out.) Therefore, the difference between the uncorrelated and correlated schemes does not seem to be in recruitment during the initial period of the release event.

We next explored whether the correlated multi-*τ* gating scheme was more efficient at terminating a Ca^2+^ release event. During a release event, the number of open channels fluctuates greatly (Fig. 2A). In order to focus on the termination phase of the event, we focused on the very end of a release event, specifically the very last time 3 RyRs were open. We wanted to gauge the speed of termination (i.e., how long it took to go from 3 RyRs open to 0) to see whether one gating scheme was more efficient than another at closing the last few channels. The multi-*τ* gating schemes were identical (46.4 ms for the uncorrelated multi-*τ* scheme and 47.1 ms for the correlated), indicating that neither is more efficient than the other at terminating an event. (Counterintuitively, the two-state scheme was much faster at 14.4 ms, despite its much longer release events. The reason for this is discussed below.)

#### Recruitment bias

To delve further into the difference between the uncorrelated and correlated multi-*τ* schemes, we focused on what occurred during the middle of the release events, the core phase of Ca^2+^ release. We reasoned that if one scheme’s Ca^2+^ release event is consistently smaller than another, then there may be a bias toward having fewer RyRs open at any one time. This can occur either by having a lower propensity to open neighboring closed RyRs or a higher propensity to close already-open RyRs. We found that the correlated gating scheme had both biases and thereby prevented Ca^2+^ release events from having a large number of open RyRs. Specifically, we computed the fraction of timesteps that *n* open RyRs became *n* +1 (Fig. 3B) or *n* −1 (Fig. 3C); that is, what is the probability that, for example, 4 RyRs open became 5 or 3?

The probability of increasing the number of open RyRs from 1 to 2 or from 2 to 3 were virtually identical, but diverged after that; there is gap between the red and blue confidence intervals in Figs. 3B and S5. Similarly, the probability of decreasing from 2 to 1 or from 1 to 0 open RyRs is the same for the two multi-*τ* gating schemes, but becomes significantly different for 3 or more open RyRs (Figs. 3C and S5). It is important to note that the small but consistent differences between the red and blue confidence intervals are statistically significant; each confidence interval was computed from 1000 bootstrap resamples of 500 100-sec-long simulations that had millions of Ca^2+^ release events (enumerated earlier), each of which had many changes in the number of open RyRs (Fig. 2A).

The net effect of the probabilities in Figs. 3B and 3C are described in Fig. 3D. There, the difference of the probabilities to increase and decrease the number of open RyRs is shown. A positive value is a bias to open more RyRs and a negative value to close RyRs. We call this the recruitment bias.

These recruitment biases are small (∼–0.005 for the uncorrelated multi-*τ* scheme when 5 RyRs are open and ∼–0.015 for the correlated multi-*τ* scheme in a 5×5 array, as shown at the maxima in the red and blue lines in Fig. 3D, respectively). However, the larger bias toward closing RyRs by the correlated multi-*τ* scheme (compared to the uncorrelated multi-*τ* scheme) has a significant effect in the Ca^2+^ release statistics toward smaller Ca^2+^ release events (i.e., fewer RyRs open per event), as seen in Figs. 2 and 3A; over the course of millions of channel openings and closings, the small recruitment bias difference has a large effect. Moreover, the ∼+0.02 bias to open more RyRs of the two-state model produces very large, long-lasting, and more frequent Ca^2+^ release events (Figs. 2 and 3A).

These results indicate that a small bias toward opening or closing RyRs during the middle of release event has substantial consequences on the global release of Ca^2+^ from the SR. They also show why the two-state model is qualitatively different from the multi-*τ* gating schemes. Specifically, it is biased toward opening more RyRs until the number of open channels becomes relatively large (black line in Fig. 3D is positive when the number of open RyRs is ≤ 9).

The fact that the two multi-*τ* gating schemes have very similar biases when just a few RyRs are open (red and blue lines in Fig. 3D) explains why both their initiation and termination statistics were nearly identical, as both of these phases of SR Ca^2+^ release involve, by definition, few open channels. During the core phase, however, many RyRs are open and so this is where the biases toward shrinking or growing the release event have an impact. Moreover, it makes sense that the correlated nature of the OTs and CTs manifests itself only during the core phase. It is only then that the number of open RyRs fluctuates in time (Fig. 2A), with RyRs opening and closing often. These channels then have previous CTs and OTs very different compared to when the RyR array is in a long quiescent state and only the occasional 1- or 2-channel release event occurs. Under these conditions, the OT/CT correlations become important and reveal themselves as a bias towards shrinking the Ca^2+^ release event, compared to the RyRs with uncorrelated OTs and CTs (the blue line in Fig. 3D is more negative than the red line).

The recruitment bias also explains the results for the two-state gating scheme in all three phases of Ca^2+^ release. During the initiation phase, the two-state scheme has a positive recruitment bias when 2 or 3 RyRs are open, unlike the negative bias of the multi-*τ* schemes. This results in significantly fewer small events being snuffed out before growing larger. Moreover, during the core phase, the positive recruitment bias when 9 or fewer RyRs are open in a 5×5 array (Fig. 3D) sustains the release event. It does this in two ways, initially by enhancing the number of open RyRs and then by delaying termination. After the negative recruitment bias for 10+ open RyRs decreases the number of open RyRs, the positive bias increases the number again and prolongs the release event. Counterintuitively, this buoying phenomenon also explains the two-state scheme’s very fast termination speed. While there is a positive recruitment bias when only a few RyRs are open, it can be overcome, but to close all remaining channels requires acting against the recruitment bias. These several low-probability RyR closings must happen in quick succession; the faster they happen, the less likely the positive recruitment bias prevents full termination. Therefore, termination is likely to be fast.

#### Composite quantities and Ca^2+^ release

Collectively, these results show that RyR group dynamics depends strongly on the details of the single-channel gating scheme, even if the RyRs have identical single-channel MOT, MCT, and *P*_o_. Therefore, none of these quantities are sufficient to predict the size and duration of Ca^2+^ release events (and therefore downstream effects like Ca^2+^ sparks). In simulations, any model of RyR gating (Markov or otherwise) must reproduce the full RyR OT and CT distributions in single-RyR simulations. Moreover, such models should also reproduce the correlations between OTs and CTs, which we show have an important role in tamping down the number of RyRs open during a Ca^2+^ release event. Since our single-RyR data were taken under cell-like conditions, these findings are likely to be physiologically relevant as well.

## CONCLUSIONS

The goal of this work was to define whether the details of RyR gating are required to predict RyR-mediated Ca^2+^ release. We did this using a very simple model of RyRs in an array. While this model is not physiological, it retains the essentials of RyRs activating each other via fluxed Ca^2+^ so that we could assess large changes in array behavior when assumptions about RyR gating were varied. Therefore, the details of RyR array behavior will change when more realistic details like Ca^2+^ buffering, non-uniform RyR array organization, and luminal SR [Ca^2+^] (20,22,33) are included. However, the qualitative differences in RyR group behavior we found should remain, as they are due to differences in single-RyR gating.

We find that measured composite (mean) quantities like *P*_o_ (or even its constituents, MOT and MCT) by themselves are insufficient to define RyR-mediated Ca^2+^ release. In fact, one important take-home message of this work is that any gating scheme should reproduce the single-channel data as faithfully as possible since that dictates what OTs and CTs the simulated channel can stochastically sample. Intuitively, if it does a poor job reproducing the OT and CT data in a simulation with only one RyR, how can one expect it to give physiologically relevant results in a simulation with many RyRs? In addition, we found that the correlations between RyR OTs and previous CTs (and vice versa) have a moderating effect on Ca^2+^ release, reducing the number of RyRs open during a release event.

Overall, this work suggests that better modeling of single-RyR2 gating will be important for better modeling of RyR2-mediated Ca^2+^ release. Previously, two-state Markov gating models were used to illustrate the dynamics of, for example, a Ca^2+^ spark. These insights have established our basic understanding of SR Ca^2+^ release. However, to model clinically-relevant physiological circumstances, more complete gating models are needed. These must go beyond reproducing mean single-RyR data but must also capture the full OT and CT distributions, the correlations between OTs and CTs, and the inability of the open RyR state to react to Ca^2+^. This will be especially important when trying to understand the effects abnormalities in RyR gating have on CRU function, for example due to proarrhythmic RyR mutations (19-22), RyR-targeted drugs or drug candidates (25-27), and diseases like atrial fibrillation and heart failure (23,24).

One way to accomplish this goal may be to move beyond Markov gating models. While they provide insights into the internal mechanisms of the RyR gating process, from a simulation point of view they are not strictly needed. Our approach that directly converts the experimental OT and CT distributions into closing and opening probabilities may be a useful alternative. It is intuitive, simple, and numerically fast, and it can be implemented in any CICR simulation.

## SUPPORTING MATERIAL

Supporting Materials includes detailed experimental and simulation methods, 5 figures, and 2 tables.

## AUTHOR CONTRIBUTIONS

DG conceived the study, wrote the code for the simulations, analyzed the data, and wrote the manuscript.

## ACKNOWLEDGMENTS

Thanks to Prof. Michael Fill for many valuable discussions and for making the experimental data available. Research reported in this publication was supported by National Heart, Lung, and Blood Institute of the National Institutes of Health under award number R01HL057832. The content is solely the responsibility of the authors and does not necessarily represent the official views of the National Institutes of Health.

## SUPPORTING REFERENCES

References 39–44 appear in the Supporting Material.

## SUPPORTING MATERIAL

## EXPERIMENTAL DETAILS

The experimental data was previously published (2) and analyzed in detail (1). Because the details are relevant for this work, here we duplicate the description from the Supporting Material of Ref. (1), updating it to include the 0.1 µM cytosolic Ca^2+^ data and omitting references to specific figures and the 100 µM luminal Ca^2+^ data not used in the present work.

Studies were undertaken with approval by the Animal Care and Use Committee of Rush University Medical Center.

Sarcoplasmic reticulum (SR) microsomes were generated from rat ventricular muscle. Microsomes were isolated as previously described (3) and stored at –80 C. Lipid bilayers (diameter 100 µm) were comprised of a 5:4:1 mixture (50 mg/ml in decane) of phosphatidylethanolamine, phosphatidylserine, and phosphatidylcholine. Solution on one side of the bilayer (cis) was virtually grounded. The cis solution initially contained a HEPES-Tris solution (250 mM HEPES and 120 mM Tris, pH 7.4). The solution on the other side of the bilayer was initially a HEPES-Ca^2+^ solution (50 mM HEPES and 10 mM Ca(OH)_2_, pH 7.4). The SR microsomes (5–15 µg) were added to the cis solution along with 500 mM CsCl and 2 mM CaCl_2_ to promote microsome fusion. Fusion of RyR2-containing microsome results in the RyR2’s cytosolic side facing the cis compartment and its luminal domains in the other compartment (4).

After single-RyR2 activity was observed, the cytosolic solution was immediately replaced to establish the various test recording conditions. The luminal solution was changed 10 minutes later. Specifically, the cytosolic recording solution contained 0.1–1000 µM of free Ca^2+^, 0.5 mM EGTA, 1 mM of free Mg^2+^, 5 mM of total ATP, 114 mM Tris, and 250 mM HEPES (pH 7.4). (All solutions were designed using the MAXC program at maxchelator.stanford.edu). The luminal recording solution contained 1000 µM free Ca^2+^ and 200 mM Cs^+^-HEPES (pH 7.4). Final recording solutions are listed in Table S1.

The 10 minute interval before changing the luminal solution means the RyR2 was exposed to 10 mM Ca^2+^, sufficiently long to promote calsequestrin (CASQ) dissociation (if any CASQ was associated with the RyR2). This CASQ stripping process is analogous to that applied by others (5-7). CSQ was stripped from the RyRs so that the RyR2 tested were not subject to CASQ-based luminal regulation and so that a homogenous population of RyRs was studied, as not all channels in this preparation are associated with CSQ (5).

All recordings were done at room temperature with current sampled at 50 µs/point (20 kHz) and filtered at 1 kHz. No correction for missing events was made. Representative current traces may be found in Ref. (2) where some of the data was previously published. The applied potential was 20, 30, or 40 mV to produce luminal-to-cytosolic cation flux. Individual recordings were performed with one applied potential, and most ionic conditions had recordings with at least two voltages. The potential did not affect *P*_o_, as shown in Fig. 1B of Ref. (1).

**Table S1.**
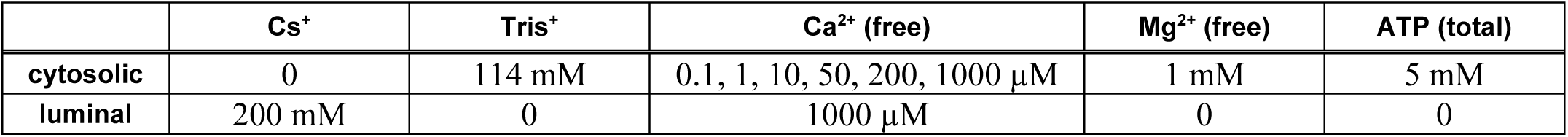
Details of the recording solutions.

Single-channel analysis was done using pCLAMP9 software (Molecular Devices). The deadtime of the filter was ∼0.185 ms. Table S2 shows details of the recordings.

**Table S2.**
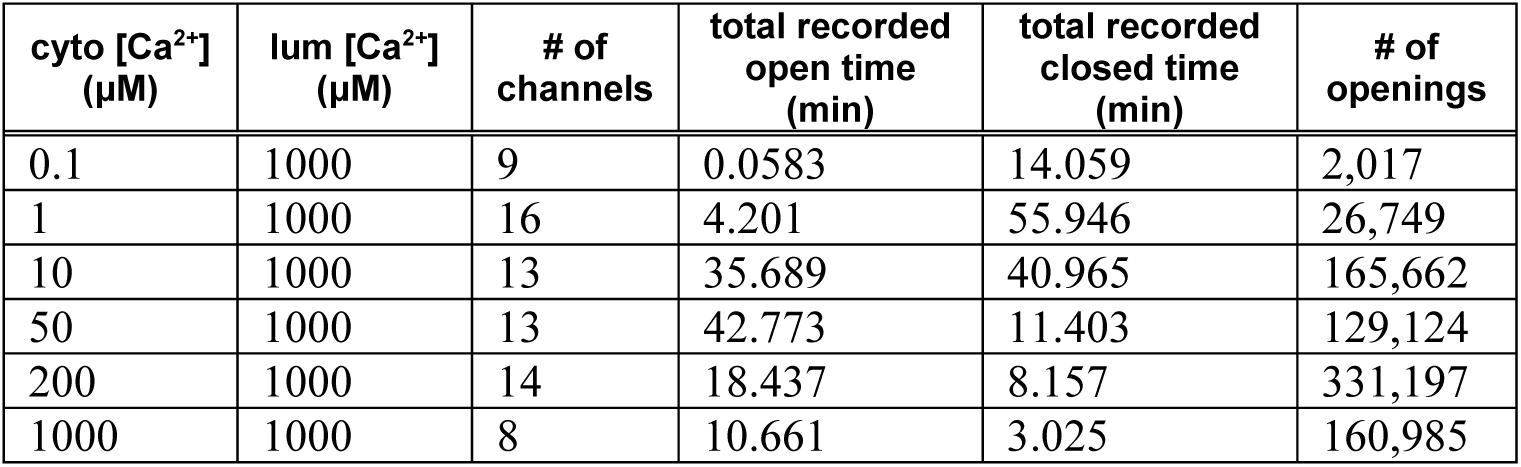
Details of the single-channel recordings: cytosolic [Ca^2+^] (column 1), luminal [Ca^2+^] (column 2), number of channels (column 3), total number of minutes in the open and closed states across all recordings (columns 4 and 5). Column 6 lists the total number of openings across all recordings, which is equal to the number of closings ±1.

## FITTING EXPONENTIALS TO OT AND CT DISTRIBUTIONS

For each cytosolic [Ca^2+^] (Table S1), open time (OT) and closed time (CT) distributions were fit by first converting them to logarithmic times and fitting the log-converted hyperexponential probability density function (pdf), as suggested by Sigworth and Sine (8). Specifically, open (subscript *o*) and closed (subscript *c*) time hyperexponential pdfs were

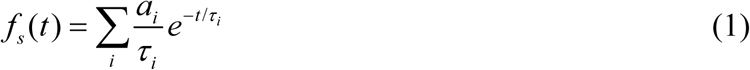

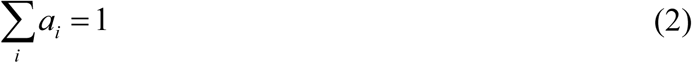

where *x* = *o* or *c*, and the log-transformed pdfs were

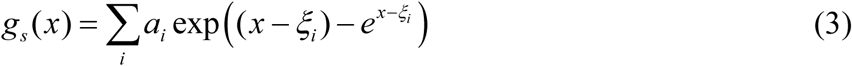

where *ξ*_*i*_ = ln(*τ*_*i*_) and *x* = ln(*t*).

Eq. (3) was fit to the log-dwell times using Mathematica 11.3 (Wolfram Research, Champaign, IL, USA) subject to the constraint that MXT (i.e., MOT or MCT) be preserved:

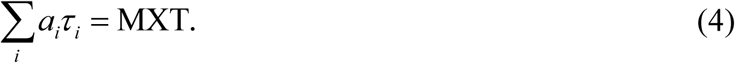

The largest time constant was determined prior to this by fitting a line to the last few points of the log-count of the log-dwell time histogram. This produced more reliable long-time fits.

The entire OT or CT data set were used (for a given cytosolic [Ca^2+^]) without excluding any events. For the two-state gating scheme, the time constant is the MXT and *a* is 1. For the uncorrelated multi-*τ* gating scheme, the data histograms used for fitting used 0.1-wide bin on the ln-time scale.

For the correlated multi-*τ* gating scheme, all consecutive pairs of closures and openings were grouped into small log-time CT bins and then these bins were combined so that there were at least 1000 openings in each bin to ensure a sufficient number of events for proper fitting. The OTs from each CT bin when then fit using the same method described for the uncorrelated multi-*τ* scheme. A similar procedure was done for pairs of consecutive openings and closures.

**Fig S1.**
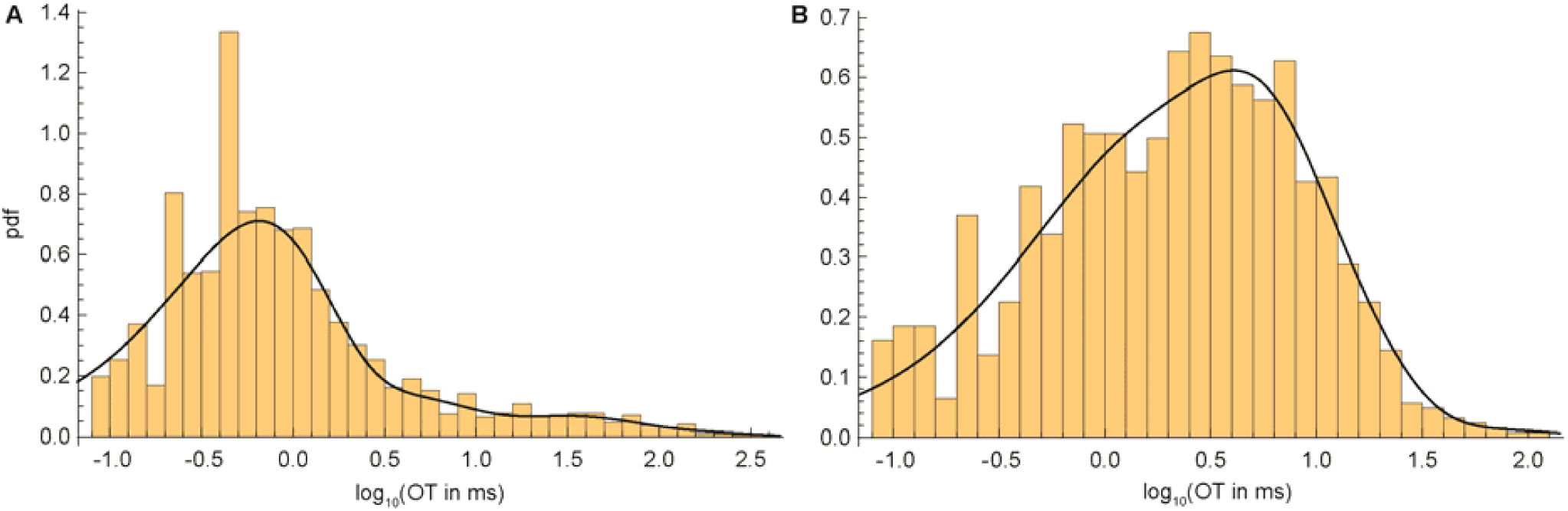
Fitted pdfs (lines) and histograms (bars) of OTs for 1 µM cytosolic [Ca^2+^] for CT (A) between 0 and 0.531 ms and (B) greater than 530.9 ms.

All fits were checked for quality by visual inspection and metrics like *R*^2^ values. Representative fits of the correlated multi-*τ* gating scheme are shown in Fig, S1. They also show why correlated OTs and CTs may give different results; the pdfs are different for short CTs preceding the opening (Fig. S1A) and long previous closures (Fig. S1B).

Our goal was solely to fit and reproduce the data as best as possible, not to determine the optimal number of exponentials that define Markov gating scheme (e.g., using maximum likelihood fitting algorithms). In fact, we do not construct any Markov schemes, and therefore it should noted that in this paper the terms “open state” and “closed state” are used exclusively to mean the conducting and nonconducting states when the current is on or off, respectively, and not the multiple “open” and “closed” states used in Markov gating models.

### Two-state model and missed events

One possible reason the two-state model may fail to more faithfully reproduce the multi-*τ* gating schemes’ results is that our experimental data was not corrected for missed events that were too short to resolve. Then the two-state Markov model may potentially generate many short events and thereby may influence (re)triggerability. We applied the two-state model correction of Roux and Sauvé (9) to test this, but only small quantitative differences (and no qualitative differences) were found between the missing-events-corrected two-state model and the uncorrected one (data not shown).

Specifically, we set the minimum time interval resolution (*τ* _*m*_) to be twice the deadtime of the filter and determined new open and closed time constants (*τ* _*o*_ and *τ* _*c*_, respectively) based on the fact that, for a two-state model, the time constant is the MXT (i.e., MOT or MCT). *τ* _*o*_ and *τ* _*c*_ are the “real” MXT (i.e., missing event corrected). Knowing the measured mean open and closed times (denoted 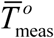 and 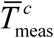, respectively), these are related by (9)

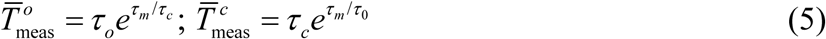

which may be solved numerically for the new time constants. These are substantially different only when 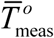 or 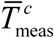 are small (e.g., MOT at low cytosolic [Ca^2+^]), but overall made not impact on RyR group dynamics.

## SIMULATION DETAILS

All simulations were performed using Mathematica 11.3 (Wolfram Research, Champaign, IL, USA) with custom-written code.

### Flow chart

At the beginning of each timestep each RyR in the array has associated with it 4 pieces of data: 1) its current state *s* (open *s* = *o* or closed *s* = *c*); 2) *T*_now_, the amount of time it has been in that state (i.e., since it last flipped states); 3) *T*_prev_, the duration of the previous state (e.g., if currently open, the length of the previous closure); 4) *C*, a cytosolic [Ca^2+^]. If the channel is closed, then *C* is the [Ca^2+^] at the center of the channel at the end of the last timestep. If, however, the channel is open, then the channel does not react to Ca^2+^ (1). Therefore, we define *C* as the [Ca^2+^] of the closed channel in the timestep before it opened.

When the simulation starts, all channels are closed and *C* is the background cytosolic [Ca^2+^]. For each channel, *T*_now_ is a random time chosen from the pdf of all experimental closed times (either the one fit with one exponential when using the two-state gating scheme or the one fit with multiple exponentials when using either multi-*τ* gating scheme). *T*_prev_ is randomly chosen similarly from the pdf of open times.

During the *n*^th^ timestep of length Δ*t*, the following steps evolve the state of the channels, with each step described in detail below:

1. The radial [Ca^2+^] profile *c*(*r, t*_*n*+1_) of each RyR is calculated using Eqs. (14) and (8). Each profile includes not only the flux of the channel if it is currently open, but also the diffusion of Ca^2+^ from any previous openings; after a channel closes, its Ca^2+^ continues to diffuse.
2. The cytosolic [Ca^2+^] on the face of each RyR is computed as the sum of all these [Ca^2+^] profiles at the centers of each RyR in the array.
3. For each channel in the array we compute *C*. If a channel is closed, then its *C* becomes the [Ca^2+^] computed in step #2. If it is open, then *C* is unchanged. Thereby, the channel only reacts to cytosolic [Ca^2+^] when it is closed and the open state is defined by the [Ca^2+^] of the previous closed state (1).
4. Based on this *C*, the appropriate *f*_*s*_ (*t*) is defined by interpolating between experimental [Ca^2+^] using Eq. (18). For the correlated gating scheme, the appropriate experimental *f*_*s*_ (*t*) were chosen based on *T*_prev_. For example, if *T*_prev_ = 0.3 ms, *s* = *o*, and *C* = 1 µM, then the *f*_*o*_ (*t*) in Fig. S1A is used. If, however, *T*_prev_ = 600 ms, then the one in Fig. S1B is used.
5. The new state and associated data of each channel in the array is computed as follows:
  a. The *f*_*s*_ (*t*) from step 4 is used in Eq. (17) with *T* = *T*_now_ to compute the probability *p* that the channel will *not* change states.
  b. A uniformly-distributed random number *r* between 0 and 1 is chosen.
  c. If *r* < 1 − *p*, then the channel state flips (i.e., *s* goes from *o* to *c* or from *c* to *o*). If not, then *s* remains unchanged.
  d. If the state remained unchanged, then *T*now is updated to *T*_now_ + Δ*t* and *T*prev is unchanged. If the state changed, then *T*_now_ is set to 0 and *T*_prev_ is set to *T*_now_ + Δ*t*.

The cycle is repeated until the end of the total simulation time, usually 100 seconds.

### Computing Ca^2+^ flux

The flux from each RyR is computed as a flux from a point source diffusing radially into infinite half-space. This has the advantage of having an analytic solution (10) that, for a constant current that turns off intermittently, is fast to compute. Here, we briefly summarize the result.

The spherically-symmetric diffusion equation from a point source is

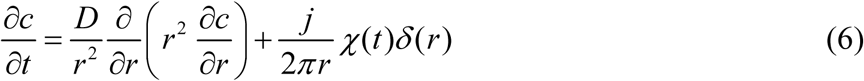

where *c*(*r, t*) is the radial (*r*) concentration profile in time (*t*) with flux *j*. *χ* (*t*) is 0 when the channel is closed and 1 when open. *δ* (*r*) is the Dirac delta-function and *D* is the diffusion coefficient. Note that we use 2*π* in the denominator of the source term instead of 4*π* because all the flux diffuses into half-space only, instead of in all radial directions; this requires doubling the current from the full radial case.

Discretizing time by *t*_*k*_ = *k*Δ*t*, we nondimensionalize by defining

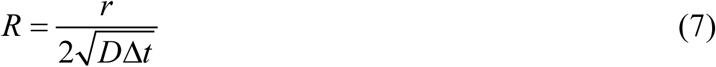

and

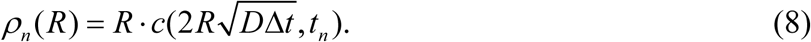

Then

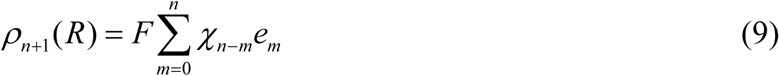

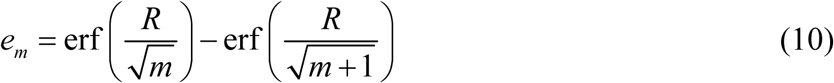

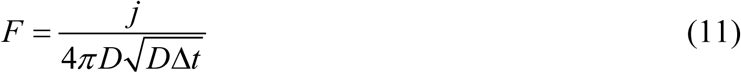

where erf is the error function and *χ*_*k*_ is 0 when the channel is closed during the time interval [*t*_*k*_, *t*_*k* +1_) and 1 when it is open. For the first term with *m* = 0, we use the relation erf(∞) = 1.

Next we take advantage of the fact that channels are open for consecutive timesteps; that is, *χ*_*k*_ = 1 for a large number of *k*. If the channel is open from *t*_*K*_ to *t*_*L*+1_, then at *t*_*n*+1_ we have

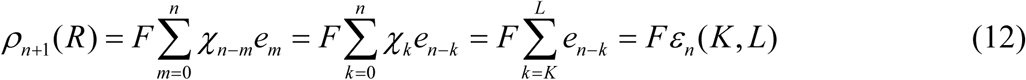

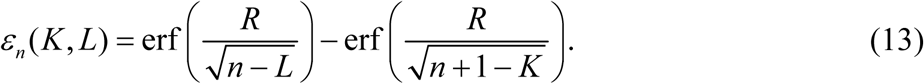

This formula is easily generalized to multiple openings (separated by a closure), indexed by *α*, when the channel is open from 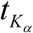 to 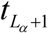:

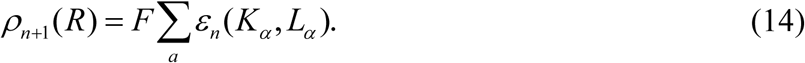

Once enough time has passed between the current time (*t*_*n*+1_) and a long-ago closure 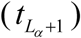, the diffused [Ca^2+^] becomes negligible and the oldest *ε*_*n*_ (*K*_*α*_, *L*_*α*_) maybe discarded (e.g., when it is 0.1% of the background [Ca^2+^]). Mathematically, *ε*_*n*_ (*K, L*) → 0 as *n* →∞.

### [Ca^2+^] at channels

The cytosolic [Ca^2+^] at each RyR in the array is the sum of currents from all channels. To avoid infinities, the [Ca^2+^] for a channel’s own flux is measured 7.5 nm away from the center while that from other channels is measured at the channel center. However, since RyRs do not respond to their own flux Ca^2+^ (they only respond to Ca^2+^ in the closed state) and Ca^2+^ diffuses quickly away from the channel when it closes, these differences in measurement location made no difference in the results.

One advantage of having a constant flux (i.e., no SR [Ca^2+^] decrease) is that when *n* channels are open during a release event, the [Ca^2+^] felt by the closed channels in the array has the same distribution no matter how many RyRs are in the array and no matter the gating scheme (data not shown). It does, of course, depend on the number of open channels *n* and the unitary RyR flux. Therefore, any differences found with the same number of open channels at one flux is not a result of different [Ca^2+^] experienced by the closed channels. The distributions are shown in Fig. S2.

**Fig S2.**
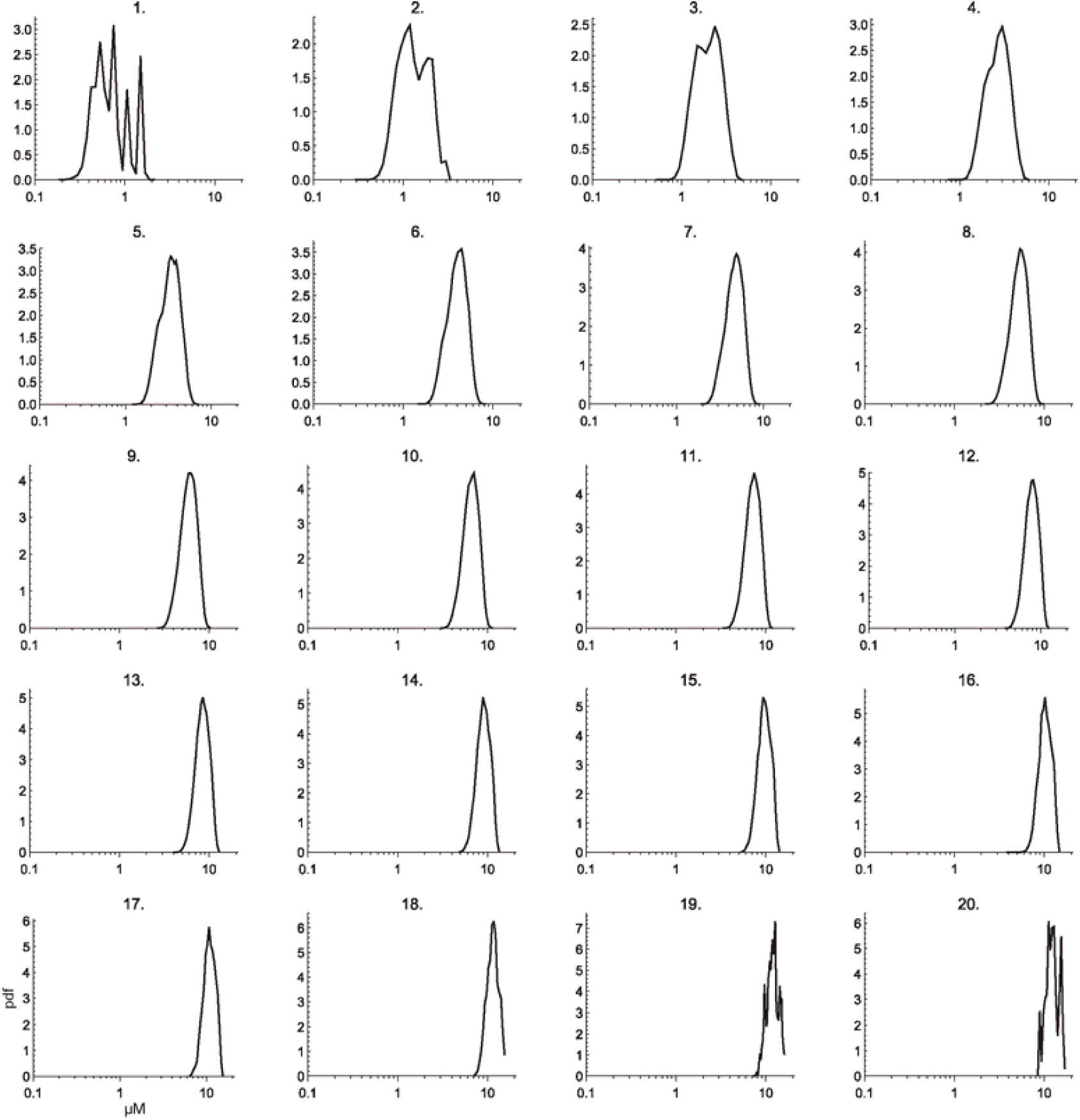
The distribution of cytosolic [Ca^2+^] felt by the closed channels in a 5×5 when *n* RyRs are open. *n* is shown at the top of each panel. The flux is 25,000.

### Stochastic state flipping

#### Computing probabilities

Suppose that a channel has been open for time *T*. The probability that it will close during the next timestep Δ*t* is the conditional probability that it will close between *T* and *T* + Δ*t* given that it has already been open for time *T* (11). Using the shorthand notation of *o* @ *t* and denote open at time *t* and closed at time *t*, respectively, this conditional probability is

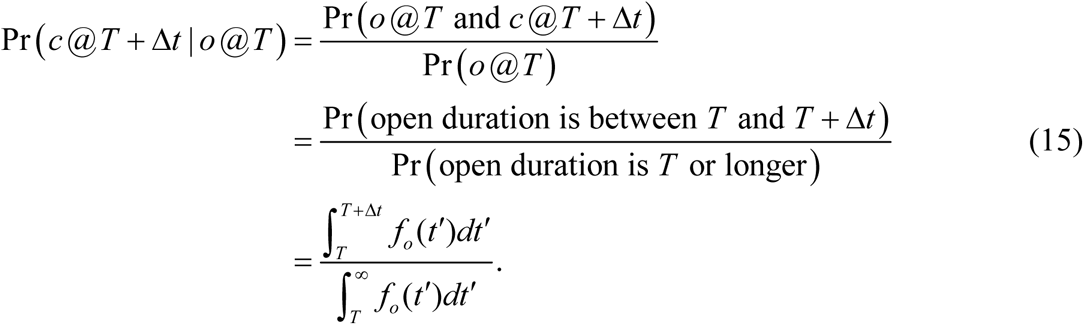

Numerically it is somewhat faster to compute the probability of staying in the same state:

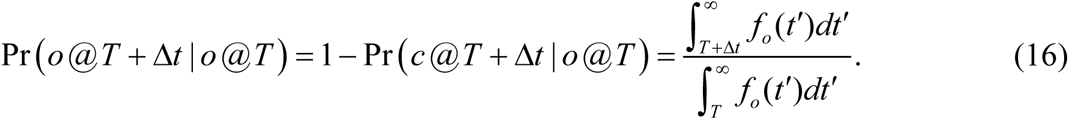

For *f*_*o*_ (*t*) given by Eq. (1), this becomes

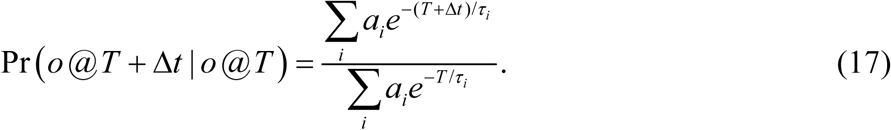

The formula for a closed channel to open is similar.

One important consequence of Eq. (17) is that it shows that the two-state case (with only a single exponential to fit the data) is qualitatively different from having multiple exponentials: the probability of flipping states is independent of *T*, the length of time the channel was already open; Eq. (17) gives that the two-state flipping probability is always 1 − *e*^−Δ*t* /*τ*^.

#### Dwell time distributions at non-experimental cytosolic [Ca^2+^]

The experiments were performed the six cytosolic [Ca^2+^] listed in Table S1. Therefore, we have the *f*_*s*_ (*t*) of Eq. (1) only for these [Ca^2+^] while the simulations will produce a continuum of [Ca^2+^]. To interpolate the *f*_*s*_ (*t*) for a specific [Ca^2+^] (i.e., the [Ca^2+^] at the center of a channel produced from neighboring open RyRs) we linearly the interpolate between the logarithm of the experimental concentrations that bracket the needed [Ca^2+^]. For example, if the needed [Ca^2+^] is *c* and the bracketing experimental concentrations are *c*_1_ and *c*_2_ and if log10 (*c*) is fraction *α* between log_10_ (*c*_1_) and log_10_ (*c*_2_), then we use

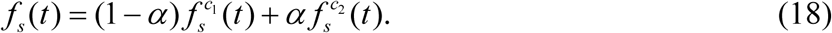

Because the individual 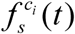 are pdfs, so is the new *f*_*s*_ (*t*).

### RyR array geometry

The RyR2s were arranged in a square arrays with center-to-center distance of 28 nm, consistent with recent experimental findings (12). The main focus was on arrays of size 5×5 and 7×7, but simulations for 3×3, 10×10, and 14×14 were also performed.

### Parameters

For all simulations we used Δ*t* = 100 µs, 0.1 µM background cytosolic [Ca^2+^] for multi-channel simulations (the stated background [Ca^2+^] for single-RyR simulations), and a Ca^2+^ diffusion coefficient of 1.58 ·10^−10^ m^2^/s. This diffusion coefficient is 20% of the experimental value to mimic the slow diffusion in the subsarcolemmal space and is similar to the value of 1.4 ·10^−10^ m^2^/s used by Cannell et al. (13).

### Correlated single-RyR simulations

The correlated multi-*τ* gating scheme reproduces the OT/CT correlations (1), as shown in Fig. S3. The two-state and uncorrelated multi-*τ* gating schemes produce flat lines (data not shown).

Experimentally, these correlations are found consistently in the presence and absence of calsequestrin and in species other than rat (unpublished data from Michael Fill, Rush University). While the origin of the correlations is unknown, they are not due to modal gating. This is shown in the data by the narrow confidence bands in the Fig. S3, meaning the open time of an event is very close to the mean open time of similar events; modal gating would produce a wider intervals due to the changing of modes between short and long openings. Moreover, the simulations show this by reproducing the experimental correlations without having modal gating in them; the gating scheme uses the experimental data as a whole, and thus does not include or produce modal gating.

**Fig S3.**
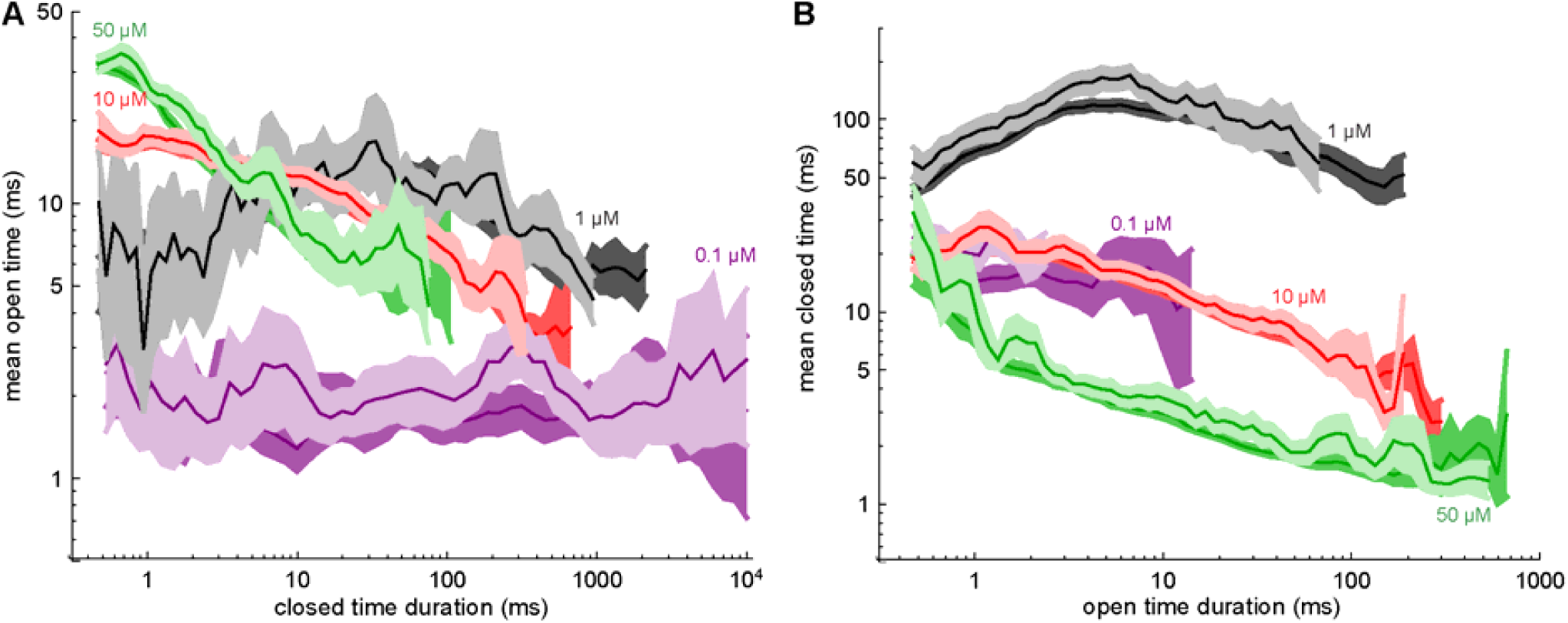
(A) Correlations between CT and the previous events’ mean OT. (B) Correlations between OT and the previous events’ mean CT. In both panels, the experimental data and 95% confidence intervals are in the dark shades and the single-channel simulation results using the correlated multi-*τ* gating scheme in the light shades. Confidence intervals were computed as described in Ref. (1) by bootstrap resampling of entire experimental and simulated records. The cytosolic [Ca^2+^] is shown for each curve. All the experimental curves except the purple 0.1 µM one are the same as in Ref. (1) except that confidence intervals are slightly different because the bootstrapping, a random resampling process, was redone here.

## RESULTS FOR DIFFERENT CLUSTER SIZES

The rate at which Ca^2+^ release events happen (i.e., their frequency) is shown in detail in Fig. S4A–C. Specifically, the frequency of release events with a maximum of 1 (left column), 2 (middle column), and 3 or more (right column) open RyRs is shown. The frequency of these latter events decreases sharply at a threshold flux because above that threshold the release events never terminate and so, in the extreme, there is only one very long event per 100-sec-long simulation. Consequently, the smaller release event frequency also drops off.

**Fig S4.**
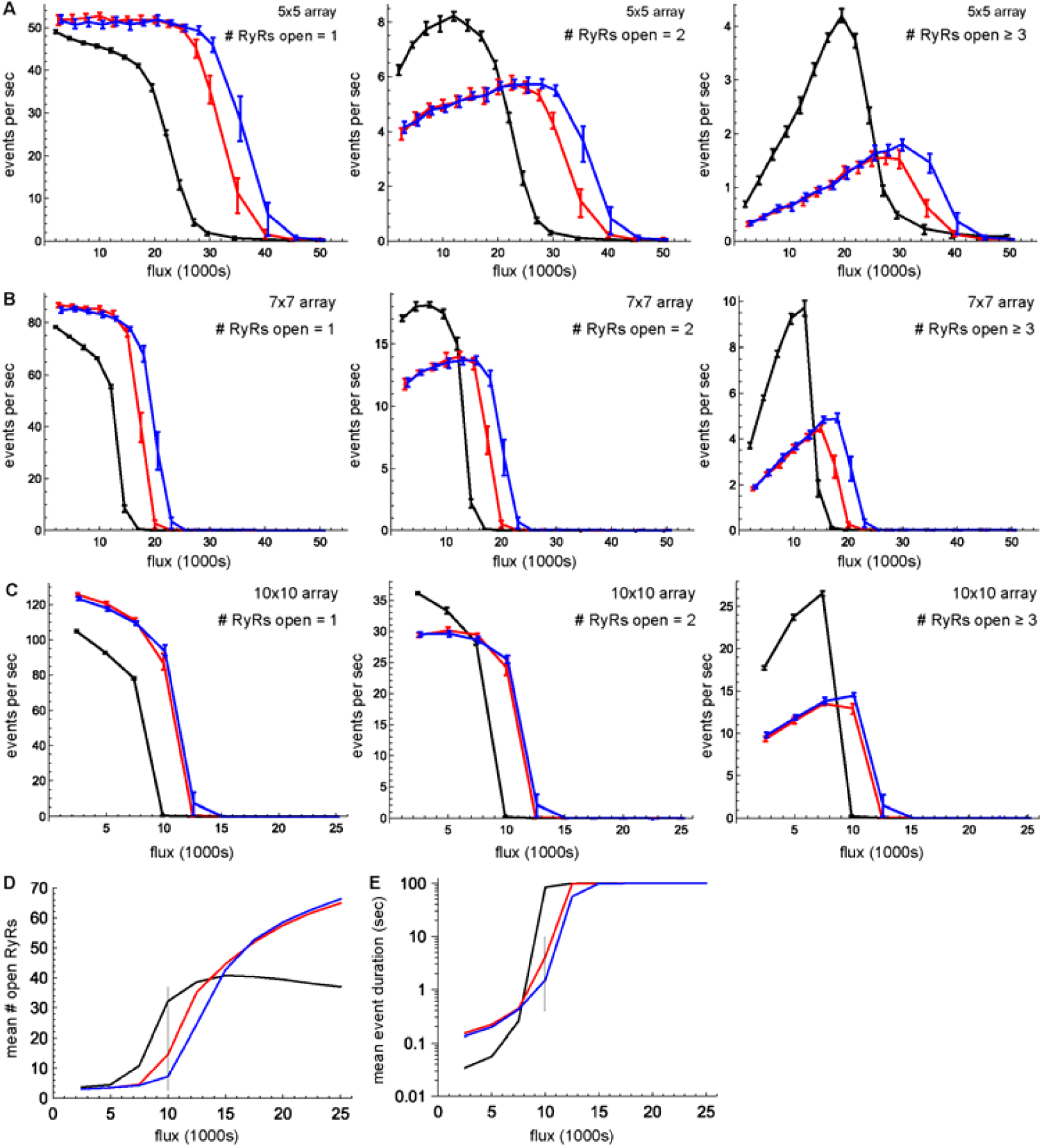
The frequency of various types of Ca^2+^ release events versus unitary RyR Ca^2+^ flux for arrays of size (A) 5×5, (B) 7×7, and (C) 10×10. Left column: only 1 RyR open during the event. Middle column: up to 2 RyRs were open. Right column: 3 or more RyRs were open. The line connects the mean of 25 separate simulations and the error bars are the 25^th^ and 75^th^ percentiles of the event frequency across those 25 simulations. Black lines: two-state gating scheme. Red lines: uncorrelated multi-*τ* scheme. Blue lines: correlated multi-*τ* scheme. Panels D and E are the same as Fig. 2B and C, respectively, but for the 10×10 array.

The results shown in Fig. 3 in the main text for the 5×5 array are shown in Fig. S5 for the 7×7 and 10×10 arrays.

**Fig S5.**
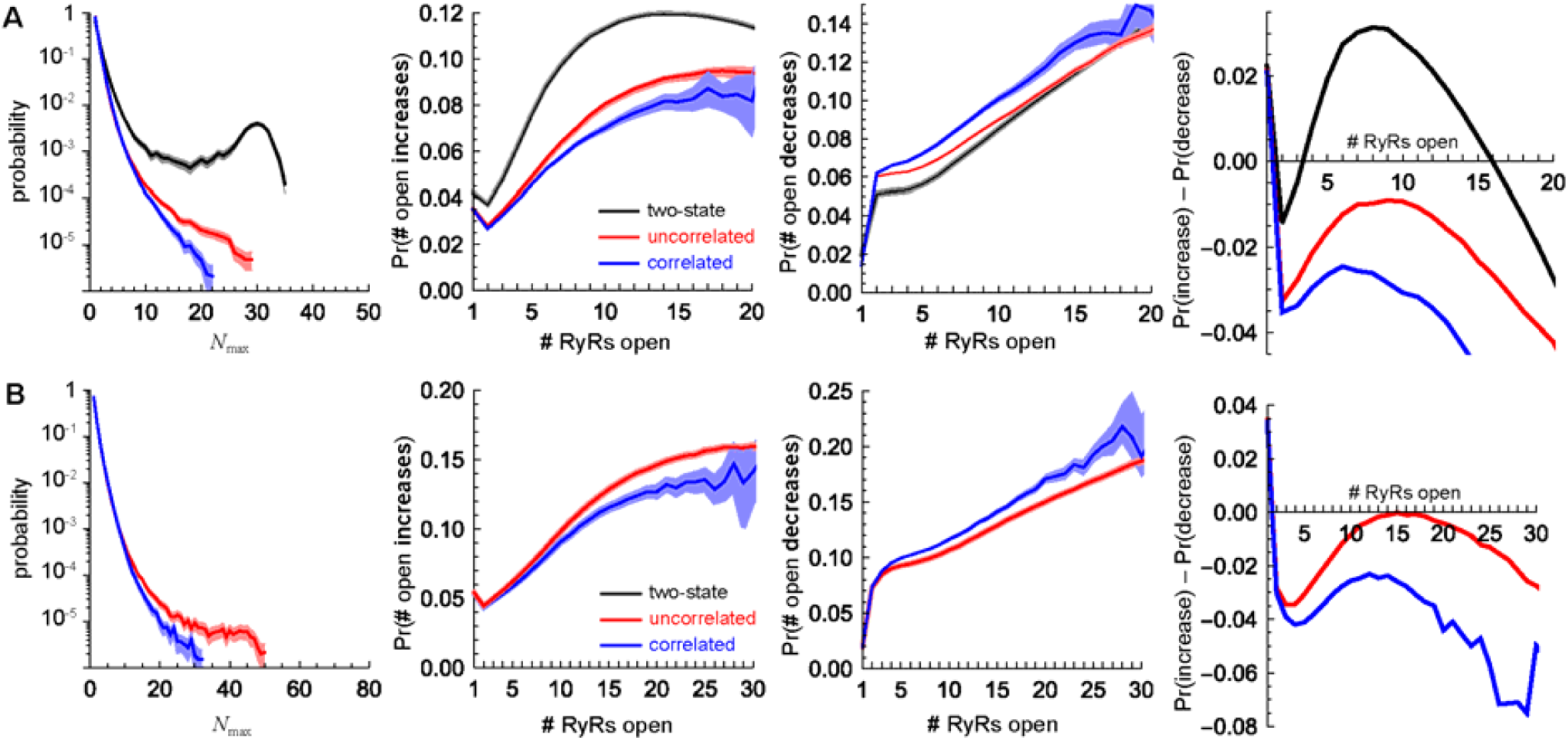
Same as Fig. 3 in the main text, but for (A) 7×7 and (B) 10×10 arrays.

## REFERENCES

1. Smith, G. D., J. E. Keizer, M. D. Stern, W. J. Lederer, and H. Cheng. 1998. A simple numerical model of calcum spark formation and detection in cardiac myocytes. Biophys. J. 75:15--32.

2. Stern, M. D., L.-S. Song, H. Cheng, J. S. K. Sham, H. T. Yang, K. R. Boheler, and E. Ríos. 1999. Local control models of cardiac excitation–contraction coupling: A possible role for allosteric interactions between ryanodine receptors. J. Gen. Physiol. 113:469–489.

3. Stern, M. D., E. Ríos, and V. A. Maltsev. 2013. Life and death of a cardiac calcium spark. J. Gen. Physiol. 142:257–274.

4. Laver, D. R., C. H. T. Kong, M. S. Imtiaz, and M. B. Cannell. 2013. Termination of calcium-induced calcium release by induction decay: An emergent property of stochastic channel gating and molecular scale architecture. J. Mol. Cell. Cardiol. 54:98–100.

5. Cannell, M. B., C. H. T. Kong, M. S. Imtiaz, and D. R. Laver. 2013. Control of sarcoplasmic reticulum Ca^2+^ release by stochastic RyR gating within a 3D model of the cardiac dyad and importance of induction decay for CICR termination. Biophys. J. 104:2149–2159.

6. Sobie, E. A., K. W. Dilly, J. dos Santos Cruz, W. J. Lederer, and M. Saleet Jafri. 2002. Termination of cardiac Ca^2+^ sparks: An investigative mathematical model of calcium-induced calcium release. Biophys. J. 83:59–78.

7. Ramay, H. R., O. Z. Liu, and E. A. Sobie. 2011. Recovery of cardiac calcium release is controlled by sarcoplasmic reticulum refilling and ryanodine receptor sensitivity. Cardiovasc. Res. 91:598–605.

8. Williams, G. S. B., A. C. Chikando, H.-T. M. Tuan, E. A. Sobie, W. J. Lederer, and M. S. Jafri. 2011. Dynamics of calcium sparks and calcium leak in the heart. Biophys. J. 101:1287–1296.

9. Wescott, A. P., M. S. Jafri, W. J. Lederer, and G. S. B. Williams. 2016. Ryanodine receptor sensitivity governs the stability and synchrony of local calcium release during cardiac excitation-contraction coupling. J. Mol. Cell. Cardiol. 92:82–92.

10. Restrepo, J. G., J. N. Weiss, and A. Karma. 2008. Calsequestrin-mediated mechanism for cellular calcium transient alternans. Biophys. J. 95:3767–3789.

11. Sato, D. and Donald M. Bers. 2011. How does stochastic ryanodine receptor-mediated Ca leak fail to initiate a Ca spark? Biophys. J. 101:2370–2379.

12. Walker, M. A., G. S. B. Williams, T. Kohl, S. E. Lehnart, M. S. Jafri, J. L. Greenstein, W. J. Lederer, and R. L. Winslow. 2014. Superresolution modeling of calcium release in the heart. Biophys. J. 107:3018–3029.

13. Gillespie, D. and M. Fill. 2013. Pernicious attrition and inter-RyR2 CICR current control in cardiac muscle. J. Mol. Cell. Cardiol. 58:53–58.

14. Zsolnay, V., M. Fill, and D. Gillespie. 2018. Sarcoplasmic reticulum Ca^2+^ release uses a cascading network of intra-SR and channel countercurrents. Biophys. J. 114:462–473.

15. Mukherjee, S., N. L. Thomas, and A. J. Williams. 2012. A mechanistic description of gating of the human cardiac ryanodine receptor in a regulated minimal environment. J. Gen. Physiol. 140:139–158.

16. Zahradník, I., S. Györke, and A. Zahradníková. 2005. Calcium activation of ryanodine receptor channels—Reconciling RyR gating models with tetrameric channel structure. J. Gen. Physiol. 126:515–527.

17. Hake, J., A. G. Edwards, Z. Yu, P. M. Kekenes-Huskey, A. P. Michailova, J. A. McCammon, M. J. Holst, M. Hoshijima, and A. D. McCulloch. 2012. Modelling cardiac calcium sparks in a three-dimensional reconstruction of a calcium release unit. J. Physiol. 590:4403–4422.

18. Colquhoun, D. and A. G. Hawkes. 1995. The principles of the stochastic interpretation of ion-channel mechanisms. In Single-Channel Recording. B. Sakmann and E. Neher, editors. 2nd ed. Plenum Press. New York. 397–482.

19. Wehrens, X. H. T., S. E. Lehnart, F. Huang, J. A. Vest, S. R. Reiken, P. J. Mohler, J. Sun, S. Guatimosim, L.-S. Song, N. Rosemblit, J. M. D’Armiento, C. Napolitano, M. Memmi, S. G. Priori, W. J. Lederer, and A. R. Marks. 2003. FKBP12.6 deficiency and defective calcium release channel (ryanodine receptor) function linked to exercise-induced sudden cardiac death. Cell 113:829–840.

20. Jiang, D., R. Wang, B. Xiao, H. Kong, D. J. Hunt, P. Choi, L. Zhang, and S. R. W. Chen. 2005. Enhanced store overload-induced Ca^2+^ release and channel sensitivity to luminal Ca^2+^ activation are common defects of RyR2 mutations linked to ventricular tachycardia and sudden death. Circ. Res. 97:1173–1181.

21. Liu, Y., J. Wei, S. M. Wong King Yuen, B. Sun, Y. Tang, R. Wang, F. Van Petegem, and S. R. W. Chen. 2017. CPVT-associated cardiac ryanodine receptor mutation G357S with reduced penetrance impairs Ca^2+^ release termination and diminishes protein expression. PLOS ONE 12:e0184177.

22. Uehara, A., T. Murayama, M. Yasukochi, M. Fill, M. Horie, T. Okamoto, Y. Matsuura, K. Uehara, T. Fujimoto, T. Sakurai, and N. Kurebayashi. 2017. Extensive Ca^2+^ leak through K4750Q cardiac ryanodine receptors caused by cytosolic and luminal Ca^2+^ hypersensitivity. J. Gen. Physiol. 149:199–218.

23. Kubalova, Z., D. Terentyev, S. Viatchenko-Karpinski, Y. Nishijima, I. Györke, R. Terentyeva, D. N. Q. da Cuñha, A. Sridhar, D. S. Feldman, R. L. Hamlin, C. A. Carnes, and S. Györke. 2005. Abnormal intrastore calcium signaling in chronic heart failure. Proc. Natl. Acad. Sci. U. S. A. 102:14104–14109.

24. Vest, J. A., X. H. T. Wehrens, S. R. Reiken, S. E. Lehnart, D. Dobrev, P. Chandra, P. Danilo, U. Ravens, M. R. Rosen, and A. R. Marks. 2005. Defective cardiac ryanodine receptor regulation during atrial fibrillation. Circulation 111:2025–2032.

25. Zhou, Q., J. Xiao, D. Jiang, R. Wang, K. Vembaiyan, A. Wang, C. D. Smith, C. Xie, W. Chen, J. Zhang, X. Tian, P. P. Jones, X. Zhong, A. Guo, H. Chen, L. Zhang, W. Zhu, D. Yang, X. Li, J. Chen, A. M. Gillis, H. J. Duff, H. Cheng, A. M. Feldman, L.-S. Song, M. Fill, T. G. Back, and S. R. W. Chen. 2011. Carvedilol and its new analogs suppress arrhythmogenic store overload-induced Ca^2+^ release. Nat. Med. 17:1003–1009.

26. Zhang, J., Q. Zhou, C. D. Smith, H. Chen, Z. Tan, B. Chen, A. Nani, G. Wu, L.-S. Song, M. Fill, T. G. Back, and S. R. W. Chen. 2015. Non-β-blocking R-carvedilol enantiomer suppresses Ca^2+^ waves and stress-induced ventricular tachyarrhythmia without lowering heart rate or blood pressure. Biochem. J. 470:233–242.

27. Tan, Z., Z. Xiao, J. Wei, J. Zhang, Q. Zhou, C. D. Smith, A. Nani, G. Wu, L.-S. Song, T. G. Back, M. Fill, and S. R. W. Chen. 2016. Nebivolol suppresses cardiac ryanodine receptor-mediated spontaneous Ca^2+^ release and catecholaminergic polymorphic ventricular tachycardia. Biochem. J. 473:4159–4172.

28. Fill, M. and D. Gillespie. 2018. Ryanodine receptor open times are determined in the closed state. Biophys. J. 115:1160–1165.

29. Maltsev, A. V., V. A. Maltsev, and M. D. Stern. 2017. Clusters of calcium release channels harness the Ising phase transition to confine their elementary intracellular signals. Proc. Natl. Acad. Sci. in press (doi: 10.1073/pnas.1701409114).

30. Maltsev, A. V., M. D. Stern, and V. A. Maltsev. 2019. Mechanisms of Calcium Leak from Cardiac Sarcoplasmic Reticulum Revealed by Statistical Mechanics. Biophys. J. 116:2212–2223.

31. Stern, M. D. 1992. Theory of excitation-contraction coupling in cardiac muscle. Biophys. J. 63:497–517.

32. Chen, H., G. Valle, S. Furlan, A. Nani, S. Gyorke, M. Fill, and P. Volpe. 2013. Mechanism of calsequestrin regulation of single cardiac ryanodine receptor in normal and pathological conditions. J. Gen. Physiol. 142:127–136.

33. Laver, D. R. 2007. Ca^2+^ stores regulate ryanodine receptor Ca^2+^ release channels via luminal and cytosolic Ca^2+^ sites. Biophys. J. 92:3541–3555.

34. Laver, D. R. and B. N. Honen. 2008. Luminal Mg^2+^, a key factor controlling RYR2-mediated Ca^2+^ release: Cytoplasmic and luminal regulation modeled in a tetrameric channel. J. Gen. Physiol. 132:429–446.

35. Shannon, T. R., F. Wang, J. Puglisi, C. Weber, and D. M. Bers. 2004. A mathematical treatment of integrated Ca dynamics within the ventricular myocyte. Biophys. J. 87:3351–3371.

36. Peskoff, A., J. A. Posit, and G. A. Langer. 1992. Sarcolemmal calcium binding sites in heart: II. Mathematical model for diffusion of calcium released from the sarcoplasmic reticulum into the diadic region. J. Membr. Biol. 129:59–69.

37. Gillespie, D. 2015. Algorithm for the time-propagation of the radial diffusion equation based on a Gaussian quadrature. PLoS ONE 10:e0132273.

38. Yeomans, J. M. 1992. Statistical Mechanics of Phase Transitions. Oxford: Oxford Univeristy Press. 153 p.

39. Chamberlain, B. K. and S. Fleischer. 1988. Isolation of canine cardiac sarcoplasmic reticulum. Methods Enzymol. 157:91–99.

40. Tu, Q., P. Velez, M. Brodwick, and M. Fill. 1994. Streaming potentials reveal a short ryanodine-sensitive selectivity filter in cardiac Ca^2+^ release channel. Biophys. J. 67:2280–2285.

41. Qin, J., G. Valle, A. Nani, A. Nori, N. Rizzi, S. G. Priori, P. Volpe, and M. Fill. 2008. Luminal Ca^2+^ regulation of single cardiac ryanodine receptors: Insights provided by calsequestrin and its mutants. J. Gen. Physiol. 131:325–334.

42. Györke, I., N. Hester, L. R. Jones, and S. Györke. 2004. The role of calsequestrin, triadin, and junctin in conferring cardiac ryanodine receptor responsiveness to luminal calcium. Biophys. J. 86:2121–2128.

43. Beard, N. A., M. G. Casarotto, L. Wei, M. Varsányi, D. R. Laver, and A. F. Dulhunty. 2005. Regulation of ryanodine receptors by calsequestrin: Effect of high luminal Ca^2+^ and phosphorylation. Biophys. J. 88:3444–3454.

44. Cabra, V., T. Murayama, and M. Samsó. 2016. Ultrastructural analysis of self-associated RyR2s. Biophys. J. 110:2651–2662.

## REFERENCES

1. Fill, M. and D. Gillespie. 2018. Ryanodine receptor open times are determined in the closed state. Biophys. J. 115:1160–1165.

2. Chen, H., G. Valle, S. Furlan, A. Nani, S. Gyorke, M. Fill, and P. Volpe. 2013. Mechanism of calsequestrin regulation of single cardiac ryanodine receptor in normal and pathological conditions. J. Gen. Physiol. 142:127–136.

3. Chamberlain, B. K. and S. Fleischer. 1988. Isolation of canine cardiac sarcoplasmic reticulum. Methods Enzymol. 157:91–99.

4. Tu, Q., P. Velez, M. Brodwick, and M. Fill. 1994. Streaming potentials reveal a short ryanodine-sensitive selectivity filter in cardiac Ca^2+^ release channel. Biophys. J. 67:2280–2285.

5. Qin, J., G. Valle, A. Nani, A. Nori, N. Rizzi, S. G. Priori, P. Volpe, and M. Fill. 2008. Luminal Ca^2+^ regulation of single cardiac ryanodine receptors: Insights provided by calsequestrin and its mutants. J. Gen. Physiol. 131:325–334.

6. Györke, I., N. Hester, L. R. Jones, and S. Györke. 2004. The role of calsequestrin, triadin, and junctin in conferring cardiac ryanodine receptor responsiveness to luminal calcium. Biophys. J. 86:2121–2128.

7. Beard, N. A., M. G. Casarotto, L. Wei, M. Varsányi, D. R. Laver, and A. F. Dulhunty. 2005. Regulation of ryanodine receptors by calsequestrin: Effect of high luminal Ca^2+^ and phosphorylation. Biophys. J. 88:3444–3454.

8. Sigworth, F. J. and S. M. Sine. 1987. Data transformations for improved display and fitting of single-channel dwell time histograms. Biophys. J. 52:1047–1054.

9. Roux, B. and R. Sauvé. 1985. A general solution to the time interval omission problem applied to single channel analysis. Biophys. J. 48:149–158.

10. Gillespie, D. 2015. Algorithm for the time-propagation of the radial diffusion equation based on a Gaussian quadrature. PLoS ONE 10:e0132273.

11. Colquhoun, D. and A. G. Hawkes. 1995. The principles of the stochastic interpretation of ion-channel mechanisms. In Single-Channel Recording. B. Sakmann and E. Neher, editors. 2nd ed. Plenum Press. New York. 397–482.

12. Cabra, V., T. Murayama, and M. Samsó. 2016. Ultrastructural analysis of self-associated RyR2s. Biophys. J. 110:2651–2662.

13. Cannell, M. B., C. H. T. Kong, M. S. Imtiaz, and D. R. Laver. 2013. Control of sarcoplasmic reticulum Ca^2+^ release by stochastic RyR gating within a 3D model of the cardiac dyad and importance of induction decay for CICR termination. Biophys. J. 104:2149–2159.

