## Supporting Materials for "Recruiting RyRs to open in a Ca^2+^ release unit: Single-RyR gating properties make RyR group dynamics"

| | $Cs^+$ | $Tris^+$ | $Ca^{2+}$ (free) | $Mg^{2+}$ (free) | ATP (total) |
|---|---|---|---|---|---|
| **cytosolic** | 0 | 114 mM | 0.1, 1, 10, 50, 200, 1000 μM | 1 mM | 5 mM |
| **luminal** | 200 mM | 0 | 1000 μM | 0 | 0 |

Table S1. Details of the recording solutions.

Single-channel analysis was done using pCLAMP9 software (Molecular Devices). The deadtime of the filter was ~0.185 ms. Table S2 shows details of the recordings.

| cyto $[Ca^{2+}]$ (μM) | lum $[Ca^{2+}]$ (μM) | # of channels | total recorded open time (min) | total recorded closed time (min) | # of openings |
|---|---|---|---|---|---|
| 0.1 | 1000 | 9 | 0.0583 | 14.059 | 2,017 |
| 1 | 1000 | 16 | 4.201 | 55.946 | 26,749 |
| 10 | 1000 | 13 | 35.689 | 40.965 | 165,662 |
| 50 | 1000 | 13 | 42.773 | 11.403 | 129,124 |
| 200 | 1000 | 14 | 18.437 | 8.157 | 331,197 |
| 1000 | 1000 | 8 | 10.661 | 3.025 | 160,985 |

Table S2. Details of the single-channel recordings: cytosolic $[Ca^{2+}]$ (column 1), luminal $[Ca^{2+}]$ (column 2), number of channels (column 3), total number of minutes in the open and closed states across all recordings (columns 4 and 5). Column 6 lists the total number of openings across all recordings, which is equal to the number of closings ±1.

### FITTING EXPONENTIALS TO OT AND CT DISTRIBUTIONS

For each cytosolic $[Ca^{2+}]$ (Table S1), open time (OT) and closed time (CT) distributions were fit by first converting them to logarithmic times and fitting the log-converted hyperexponential probability density function (pdf), as suggested by Sigworth and Sine (8). Specifically, open (subscript *o*) and closed (subscript *c*) time hyperexponential pdfs were

$$f_s(t) = \sum_i \frac{a_i}{\tau_i} e^{-t/\tau_i} \tag{1}$$

$$\sum_i a_i = 1 \tag{2}$$

where $x = o$ or *c*, and the log-transformed pdfs were

$$g_s(x) = \sum_i a_i \exp\left((x - \xi_i) - e^{x-\xi_i}\right) \tag{3}$$

where $\xi_i = \ln(\tau_i)$ and $x = \ln(t)$.

Eq. (3) was fit to the log-dwell times using Mathematica 11.3 (Wolfram Research, Champaign, IL, USA) subject to the constraint that MXT (i.e., MOT or MCT) be preserved:

$$\sum_i a_i \tau_i = \text{MXT}. \tag{4}$$

The largest time constant was determined prior to this by fitting a line to the last few points of the log-count of the log-dwell time histogram. This produced more reliable long-time fits.

The entire OT or CT data set were used (for a given cytosolic $[Ca^{2+}]$) without excluding any events. For the two-state gating scheme, the time constant is the MXT and *a* is 1. For the uncorrelated multi-$\tau$ gating scheme, the data histograms used for fitting used 0.1-wide bin on the ln-time scale.

For the correlated multi-$\tau$ gating scheme, all consecutive pairs of closures and openings were grouped into small log-time CT bins and then these bins were combined so that there were at least 1000 openings in each bin to ensure a sufficient number of events for proper fitting. The OTs from each CT bin when then fit using the same method described for the uncorrelated multi-$\tau$ scheme. A similar procedure was done for pairs of consecutive openings and closures.

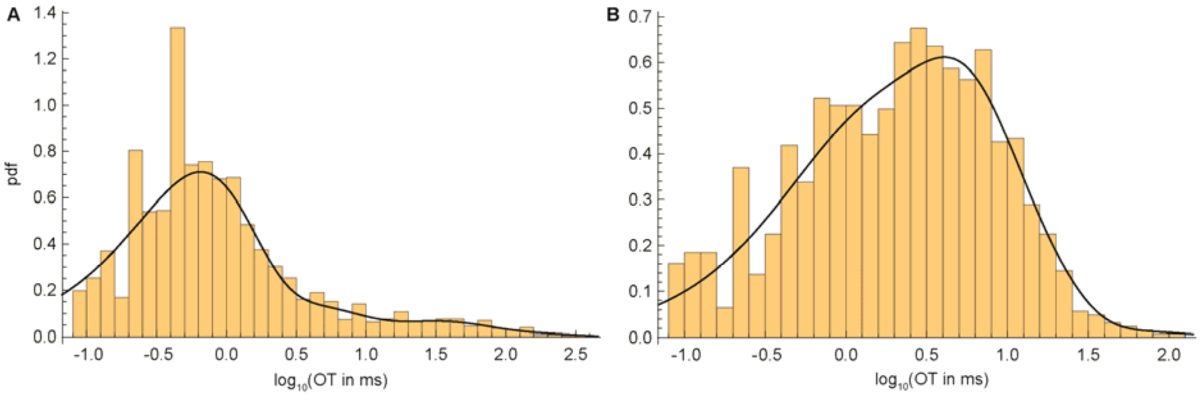


Fig. S1. Fitted pdfs (lines) and histograms (bars) of OTs for 1 μM cytosolic [$Ca^{2+}$] for CT (A) between 0 and 0.531 ms and (B) greater than 530.9 ms.

All fits were checked for quality by visual inspection and metrics like $R^2$ values. Representative fits of the correlated multi-$\tau$ gating scheme are shown in Fig, S1. They also show why correlated OTs and CTs may give different results; the pdfs are different for short CTs preceding the opening (Fig. S1A) and long previous closures (Fig. S1B).

Our goal was solely to fit and reproduce the data as best as possible, not to determine the optimal number of exponentials that define Markov gating scheme (e.g., using maximum likelihood fitting algorithms). In fact, we do not construct any Markov schemes, and therefore it should noted that in this paper the terms “open state” and “closed state” are used exclusively to mean the conducting and nonconducting states when the current is on or off, respectively, and not the multiple “open” and “closed” states used in Markov gating models.

#### *Two-state model and missed events*

One possible reason the two-state model may fail to more faithfully reproduce the multi-τ gating schemes’ results is that our experimental data was not corrected for missed events that were too short to resolve. Then the two-state Markov model may potentially generate many short events and thereby may influence (re)triggerability. We applied the two-state model correction of Roux and Sauvé (9) to test this, but only small quantitative differences (and no qualitative differences) were found between the missing-events-corrected two-state model and the uncorrected one (data not shown).

Specifically, we set the minimum time interval resolution ($\tau_m$) to be twice the deadtime of the filter and determined new open and closed time constants ($\tau_o$ and $\tau_c$, respectively) based on the fact that, for a two-state model, the time constant is the MXT (i.e., MOT or MCT). $\tau_o$ and $\tau_c$ are the “real” MXT (i.e., missing event corrected). Knowing the measured mean open and closed times (denoted $\overline{T}^o_{\text{meas}}$ and $\overline{T}^c_{\text{meas}}$, respectively), these are related by (9)

$$\overline{T}^o_{\text{meas}} = \tau_o e^{\tau_m/\tau_c}; \; \overline{T}^c_{\text{meas}} = \tau_c e^{\tau_m/\tau_0} \quad (5)$$

which may be solved numerically for the new time constants. These are substantially different only when $\overline{T}^o_{\text{meas}}$ or $\overline{T}^c_{\text{meas}}$ are small (e.g., MOT at low cytosolic [$Ca^{2+}$]), but overall made not impact on RyR group dynamics.

### SIMULATION DETAILS

All simulations were performed using Mathematica 11.3 (Wolfram Research, Champaign, IL, USA) with custom-written code.

#### Flow chart

At the beginning of each timestep each RyR in the array has associated with it 4 pieces of data: 1) its current state $s$ (open $s = o$ or closed $s = c$); 2) $T_{\text{now}}$, the amount of time it has been in that state (i.e., since it last flipped states); 3) $T_{\text{prev}}$, the duration of the previous state (e.g., if currently open, the length of the previous closure); 4) $C$, a cytosolic [$Ca^{2+}$]. If the channel is closed, then $C$ is the [$Ca^{2+}$] at the center of the channel at the end of the last timestep. If, however, the channel is open, then the channel does not react to $Ca^{2+}$ (1). Therefore, we define $C$ as the [$Ca^{2+}$] of the closed channel in the timestep before it opened.

When the simulation starts, all channels are closed and $C$ is the background cytosolic [$Ca^{2+}$]. For each channel, $T_{\text{now}}$ is a random time chosen from the pdf of all experimental closed times (either the one fit with one exponential when using the two-state gating scheme or the one fit with multiple exponentials when using either multi-$\tau$ gating scheme). $T_{\text{prev}}$ is randomly chosen similarly from the pdf of open times.

The spherically-symmetric diffusion equation from a point source is

$$\frac{\partial c}{\partial t} = \frac{D}{r^2}\frac{\partial}{\partial r}\left(r^2 \frac{\partial c}{\partial r}\right) + \frac{j}{2\pi r}\chi(t)\delta(r) \tag{6}$$

where $c(r,t)$ is the radial ($r$) concentration profile in time ($t$) with flux $j$. $\chi(t)$ is 0 when the channel is closed and 1 when open. $\delta(r)$ is the Dirac delta-function and $D$ is the diffusion coefficient. Note that we use $2\pi$ in the denominator of the source term instead of $4\pi$ because all the flux diffuses into half-space only, instead of in all radial directions; this requires doubling the current from the full radial case.

Discretizing time by $t_k = k\Delta t$, we nondimensionalize by defining

$$R = \frac{r}{2\sqrt{D\Delta t}} \tag{7}$$

and

$$\rho_n(R) = R \cdot c(2R\sqrt{D\Delta t}, t_n). \tag{8}$$

Then

$$\rho_{n+1}(R) = F\sum_{m=0}^{n} \chi_{n-m} e_m \tag{9}$$

$$e_m = \operatorname{erf}\left(\frac{R}{\sqrt{m}}\right) - \operatorname{erf}\left(\frac{R}{\sqrt{m+1}}\right) \tag{10}$$

$$F = \frac{j}{4\pi D\sqrt{D\Delta t}} \tag{11}$$

where erf is the error function and $\chi_k$ is 0 when the channel is closed during the time interval $[t_k, t_{k+1})$ and 1 when it is open. For the first term with $m = 0$, we use the relation $\operatorname{erf}(\infty) = 1$.

Next we take advantage of the fact that channels are open for consecutive timesteps; that is, $\chi_k = 1$ for a large number of $k$. If the channel is open from $t_K$ to $t_{L+1}$, then at $t_{n+1}$ we have

$$\rho_{n+1}(R) = F\sum_{m=0}^{n} \chi_{n-m} e_m = F\sum_{k=0}^{n} \chi_k e_{n-k} = F\sum_{k=K}^{L} e_{n-k} = F\varepsilon_n(K,L) \tag{12}$$

$$\varepsilon_n(K,L) = \operatorname{erf}\left(\frac{R}{\sqrt{n-L}}\right) - \operatorname{erf}\left(\frac{R}{\sqrt{n+1-K}}\right). \tag{13}$$

This formula is easily generalized to multiple openings (separated by a closure), indexed by $\alpha$, when the channel is open from $t_{K_\alpha}$ to $t_{L_\alpha+1}$:

$$\rho_{n+1}(R) = F\sum_{a} \varepsilon_n(K_\alpha, L_\alpha). \tag{14}$$

Once enough time has passed between the current time ($t_{n+1}$) and a long-ago closure ($t_{L_\alpha+1}$), the diffused [$Ca^{2+}$] becomes negligible and the oldest $\varepsilon_n(K_\alpha, L_\alpha)$ maybe discarded (e.g., when it is 0.1% of the background [$Ca^{2+}$]). Mathematically, $\varepsilon_n(K,L) \to 0$ as $n \to \infty$.

#### [$Ca^{2+}$] at channels

The cytosolic [$Ca^{2+}$] at each RyR in the array is the sum of currents from all channels. To avoid infinities, the [$Ca^{2+}$] for a channel's own flux is measured 7.5 nm away from the center while that from other channels is measured at the channel center. However, since RyRs do not respond to their own flux $Ca^{2+}$ (they only respond to $Ca^{2+}$ in the closed state) and $Ca^{2+}$ diffuses quickly away from the channel when it closes, these differences in measurement location made no difference in the results.

One advantage of having a constant flux (i.e., no SR [$Ca^{2+}$] decrease) is that when *n* channels are open during a release event, the [$Ca^{2+}$] felt by the closed channels in the array has the same distribution no matter how many RyRs are in the array and no matter the gating scheme (data not shown). It does, of course, depend on the number of open channels *n* and the unitary RyR flux. Therefore, any differences found with the same number of open channels at one flux is not a result of different [$Ca^{2+}$] experienced by the closed channels. The distributions are shown in Fig. S2.

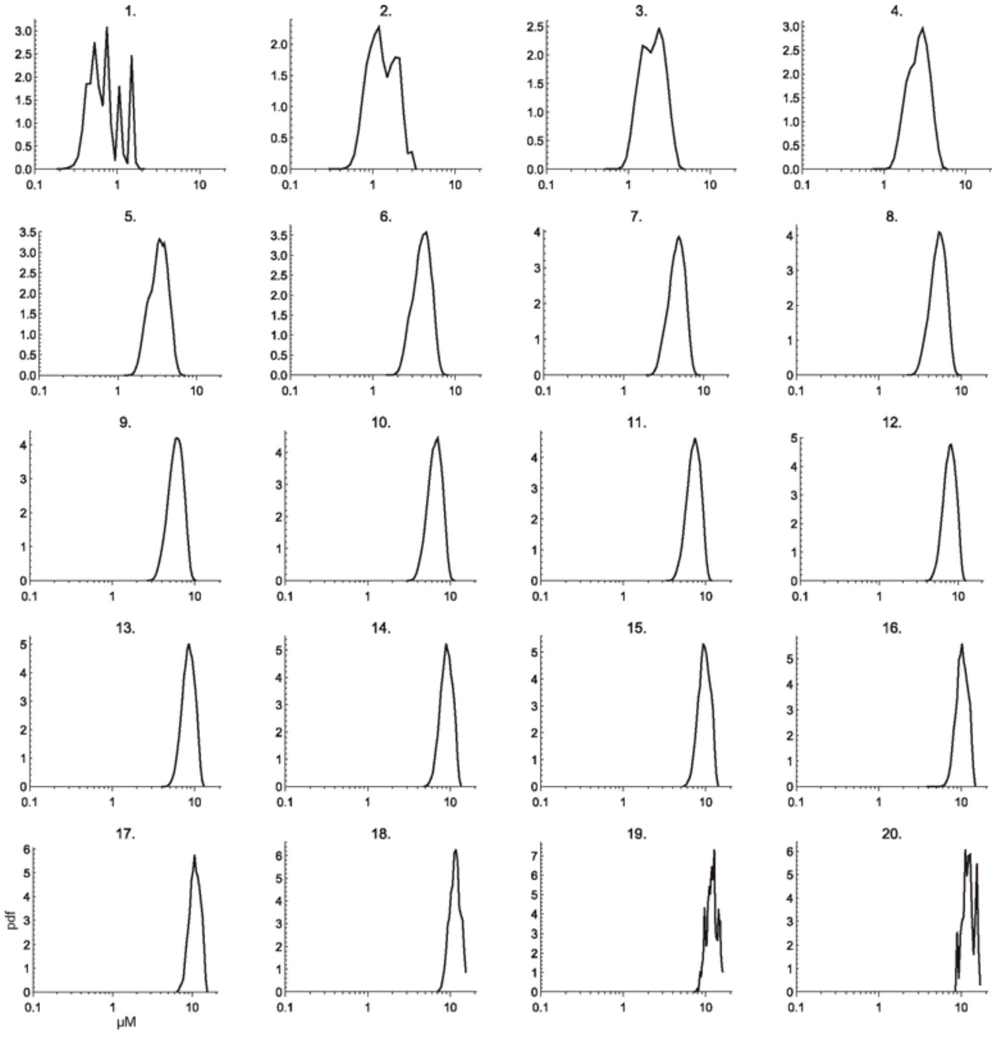


Fig. S2. The distribution of cytosolic [$Ca^{2+}$] felt by the closed channels in a 5×5 when *n* RyRs are open. *n* is shown at the top of each panel. The flux is 25,000.

#### *Stochastic state flipping*

##### Computing probabilities

Suppose that a channel has been open for time *T*. The probability that it will close during the next timestep $\Delta t$ is the conditional probability that it will close between *T* and $T+\Delta t$ given that it has already been open for time *T* (11). Using the shorthand notation of $o@t$ and $c@t$ to denote open at time *t* and closed at time *t*, respectively, this conditional probability is

$$\begin{aligned}\Pr\left(c@T+\Delta t \mid o@T\right) &= \frac{\Pr\left(o@T \text{ and } c@T+\Delta t\right)}{\Pr\left(o@T\right)} \\ &= \frac{\Pr\left(\text{open duration is between } T \text{ and } T+\Delta t\right)}{\Pr\left(\text{open duration is } T \text{ or longer}\right)} \\ &= \frac{\int_T^{T+\Delta t} f_o(t')dt'}{\int_T^{\infty} f_o(t')dt'}.\end{aligned} \tag{15}$$

Numerically it is somewhat faster to compute the probability of staying in the same state:

$$\Pr\left(o@T+\Delta t \mid o@T\right) = 1-\Pr\left(c@T+\Delta t \mid o@T\right) = \frac{\int_{T+\Delta t}^{\infty} f_o(t')dt'}{\int_T^{\infty} f_o(t')dt'}. \tag{16}$$

For $f_o(t)$ given by Eq. (1), this becomes

$$\Pr\left(o@T+\Delta t \mid o@T\right) = \frac{\sum_i a_i e^{-(T+\Delta t)/\tau_i}}{\sum_i a_i e^{-T/\tau_i}}. \tag{17}$$

The formula for a closed channel to open is similar.

One important consequence of Eq. (17) is that it shows that the two-state case (with only a single exponential to fit the data) is qualitatively different from having multiple exponentials: the probability of flipping states is independent of *T*, the length of time the channel was already open; Eq. (17) gives that the two-state flipping probability is always $1-e^{-\Delta t/\tau}$.

##### Dwell time distributions at non-experimental cytosolic [$Ca^{2+}$]

The experiments were performed the six cytosolic [$Ca^{2+}$] listed in Table S1. Therefore, we have the $f_s(t)$ of Eq. (1) only for these [$Ca^{2+}$] while the simulations will produce a continuum of [$Ca^{2+}$]. To interpolate the $f_s(t)$ for a specific [$Ca^{2+}$] (i.e., the [$Ca^{2+}$] at the center of a channel produced from neighboring open RyRs) we linearly the interpolate between the logarithm of the experimental concentrations that bracket the needed [$Ca^{2+}$]. For example, if the needed [$Ca^{2+}$] is *c* and the bracketing experimental concentrations are $c_1$ and $c_2$ and if $\log_{10}(c)$ is fraction $\alpha$ between $\log_{10}(c_1)$ and $\log_{10}(c_2)$, then we use

$$f_s(t) = (1-\alpha) f_s^{c_1}(t) + \alpha f_s^{c_2}(t). \tag{18}$$

Because the individual $f_s^{c_i}(t)$ are pdfs, so is the new $f_s(t)$.

#### RyR array geometry

The RyR2s were arranged in a square arrays with center-to-center distance of 28 nm, consistent with recent experimental findings (12). The main focus was on arrays of size 5×5 and 7×7, but simulations for 3×3, 10×10, and 14×14 were also performed.

#### Parameters

For all simulations we used $\Delta t = 100$ µs, 0.1 µM background cytosolic [$Ca^{2+}$] for multi-channel simulations (the stated background [$Ca^{2+}$] for single-RyR simulations), and a $Ca^{2+}$ diffusion coefficient of $1.58 \cdot 10^{-10}$ m$^2$/s. This diffusion coefficient is 20% of the experimental value to mimic the slow diffusion in the subsarcolemmal space and is similar to the value of $1.4 \cdot 10^{-10}$ m$^2$/s used by Cannell et al. (13).

#### Correlated single-RyR simulations

The correlated multi-$\tau$ gating scheme reproduces the OT/CT correlations (1), as shown in Fig. S3. The two-state and uncorrelated multi-$\tau$ gating schemes produce flat lines (data not shown).

Experimentally, these correlations are found consistently in the presence and absence of calsequestrin and in species other than rat (unpublished data from Michael Fill, Rush University). While the origin of the correlations is unknown, they are not due to modal gating. This is shown in the data by the narrow confidence bands in the Fig. S3, meaning the open time of an event is very close to the mean open time of similar events; modal gating would produce a wider intervals due to the changing of modes between short and long openings. Moreover, the simulations show this by reproducing the experimental correlations without having modal gating in them; the gating scheme uses the experimental data as a whole, and thus does not include or produce modal gating.

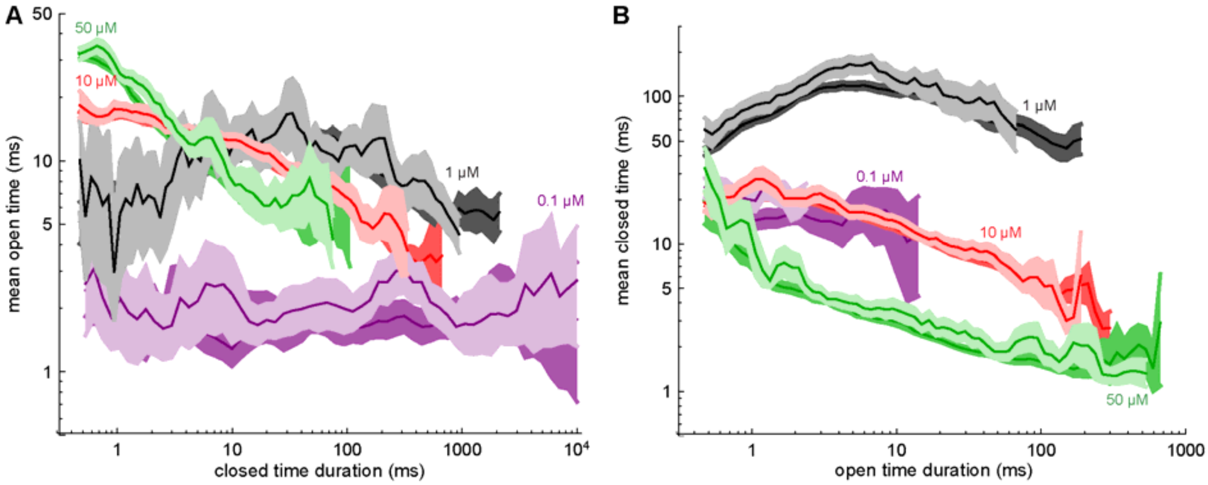


Fig. S3. (A) Correlations between CT and the previous events' mean OT. (B) Correlations between OT and the previous events' mean CT. In both panels, the experimental data and 95% confidence intervals are in the dark shades and the single-channel simulation results using the correlated multi-$\tau$ gating scheme in the light shades. Confidence intervals were computed as described in Ref. (1) by bootstrap resampling of entire experimental and simulated records. The cytosolic [$Ca^{2+}$] is shown for each curve. All the experimental curves except the purple 0.1 µM one are the same as in Ref. (1) except that confidence intervals are slightly different because the bootstrapping, a random resampling process, was redone here.

### RESULTS FOR DIFFERENT CLUSTER SIZES

The rate at which $Ca^{2+}$ release events happen (i.e., their frequency) is shown in detail in Fig. S4A–C. Specifically, the frequency of release events with a maximum of 1 (left column), 2 (middle column), and 3 or more (right column) open RyRs is shown. The frequency of these latter events decreases sharply at a threshold flux because above that threshold the release events never terminate and so, in the extreme, there is only one very long event per 100-sec-long simulation. Consequently, the smaller release event frequency also drops off.

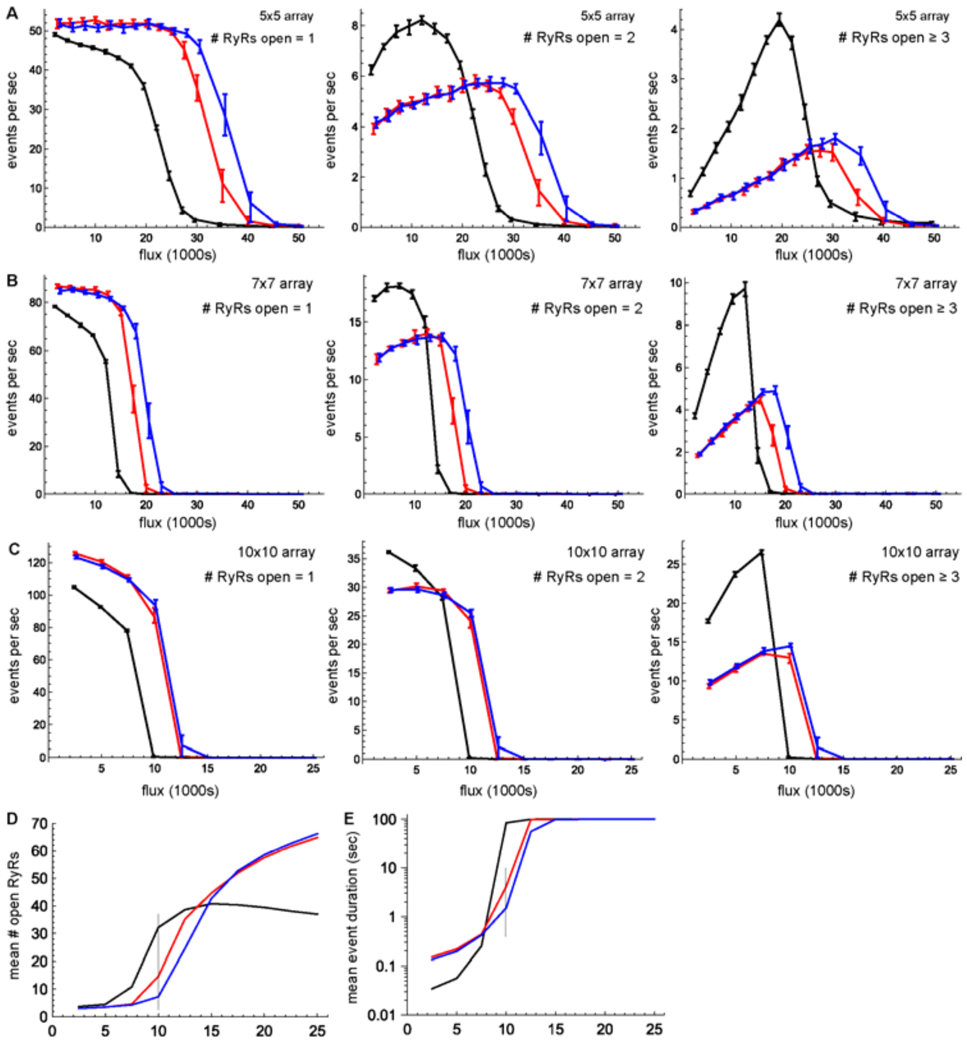


Fig. S4. The frequency of various types of $Ca^{2+}$ release events versus unitary RyR $Ca^{2+}$ flux for arrays of size (A) 5×5, (B) 7×7, and (C) 10×10. Left column: only 1 RyR open during the event. Middle column: up to 2 RyRs were open. Right column: 3 or more RyRs were open. The line connects the mean of 25 separate simulations and the error bars are the 25th and 75th percentiles of the event frequency across those 25 simulations. Black lines: two-state gating scheme. Red lines: uncorrelated multi-$\tau$ scheme. Blue lines: correlated multi-$\tau$ scheme. Panels D and E are the same as Fig. 2B and C, respectively, but for the 10×10 array.

The results shown in Fig. 3 in the main text for the 5×5 array are shown in Fig. S5 for the 7×7 and 10×10 arrays.

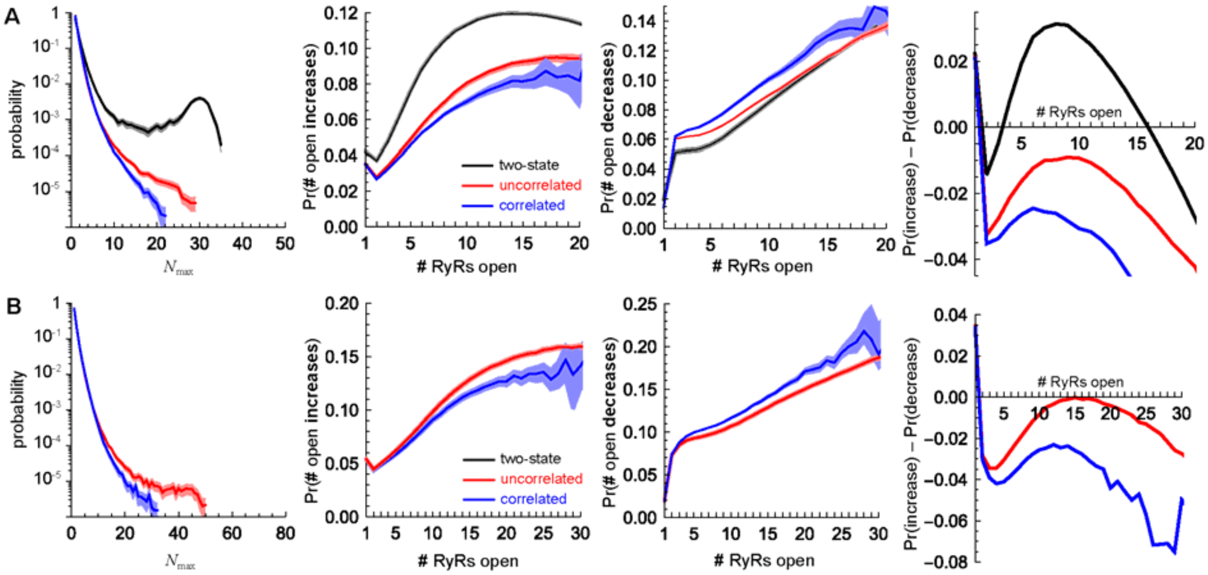


Fig. S5. Same as Fig. 3 in the main text, but for (A) 7×7 and (B) 10×10 arrays.

### REFERENCES


1. Fill, M. and D. Gillespie. 2018. Ryanodine receptor open times are determined in the closed state. *Biophys. J.* 115:1160-1165.
2. Chen, H., G. Valle, S. Furlan, A. Nani, S. Gyorke, M. Fill, and P. Volpe. 2013. Mechanism of calsequestrin regulation of single cardiac ryanodine receptor in normal and pathological conditions. *J. Gen. Physiol.* 142:127-136.
3. Chamberlain, B. K. and S. Fleischer. 1988. Isolation of canine cardiac sarcoplasmic reticulum. *Methods Enzymol.* 157:91-99.
4. Tu, Q., P. Velez, M. Brodwick, and M. Fill. 1994. Streaming potentials reveal a short ryanodine-sensitive selectivity filter in cardiac $Ca^{2+}$ release channel. *Biophys. J.* 67:2280-2285.
5. Qin, J., G. Valle, A. Nani, A. Nori, N. Rizzi, S. G. Priori, P. Volpe, and M. Fill. 2008. Luminal $Ca^{2+}$ regulation of single cardiac ryanodine receptors: Insights provided by calsequestrin and its mutants. *J. Gen. Physiol.* 131:325-334.
6. Györke, I., N. Hester, L. R. Jones, and S. Györke. 2004. The role of calsequestrin, triadin, and junctin in conferring cardiac ryanodine receptor responsiveness to luminal calcium. *Biophys. J.* 86:2121-2128.
7. Beard, N. A., M. G. Casarotto, L. Wei, M. Varsányi, D. R. Laver, and A. F. Dulhunty. 2005. Regulation of ryanodine receptors by calsequestrin: Effect of high luminal $Ca^{2+}$ and phosphorylation. *Biophys. J.* 88:3444-3454.
8. Sigworth, F. J. and S. M. Sine. 1987. Data transformations for improved display and fitting of single-channel dwell time histograms. *Biophys. J.* 52:1047-1054.
9. Roux, B. and R. Sauvé. 1985. A general solution to the time interval omission problem applied to single channel analysis. *Biophys. J.* 48:149-158.
10. Gillespie, D. 2015. Algorithm for the time-propagation of the radial diffusion equation based on a Gaussian quadrature. *PLoS ONE* 10:e0132273.
11. Colquhoun, D. and A. G. Hawkes. 1995. The principles of the stochastic interpretation of ion-channel mechanisms. In Single-Channel Recording. B. Sakmann and E. Neher, editors. 2nd ed. Plenum Press. New York. 397-482.
12. Cabra, V., T. Murayama, and M. Samsó. 2016. Ultrastructural analysis of self-associated RyR2s. *Biophys. J.* 110:2651-2662.
13. Cannell, M. B., C. H. T. Kong, M. S. Imtiaz, and D. R. Laver. 2013. Control of sarcoplasmic reticulum $Ca^{2+}$ release by stochastic RyR gating within a 3D model of the cardiac dyad and importance of induction decay for CICR termination. *Biophys. J.* 104:2149-2159.
